# Optogenetic control of PLC-γ1 activity directs cell motility

**DOI:** 10.1101/2025.10.09.681531

**Authors:** Ravikanth Appalabhotla, Priscila F. Siesser, Harrison Truscott, Nicole Hajicek, John Sondek, James E. Bear, Jason M. Haugh

## Abstract

Phospholipase C-γ1 (PLC-γ1) signaling is required for mesenchymal chemotaxis, but is it sufficient to bias motility? PLC-γ1 enzyme activity is basally autoinhibited, and light-controlled membrane recruitment of wild-type PLC-γ1 (OptoPLC-γ1) in *Plcg1-*null fibroblasts does not trigger lipid hydrolysis, complicating efforts to isolate its contribution. Utilizing cancer-associated mutations to investigate the regulatory logic of PLC-γ1, we demonstrate that a hallmark of enzyme activity, phosphorylated Tyr783 (pTyr783), is not a proxy for activity level, but is rather a marker of dysregulated autoinhibition. Accordingly, OptoPLC-γ1 with a deregulating mutation (P867R, S345F, or D1165H) exhibits elevated phosphorylation, and membrane localization of such is sufficient to activate substrate hydrolysis and concomitant motility responses. In particular, local recruitment of OptoPLC-γ1 S345F polarizes cell motility and migration on demand. This response is spatially dose-sensitive and only partially reduced by blocking canonical PLC-γ1 signaling yet is lipase-dependent. Our findings reframe the interpretation of PLC-γ1 regulation and demonstrate that local activation of PLC-γ1 is sufficient to direct cell motility.

## INTRODUCTION

Directed cell migration—a process in which cells navigate using spatiotemporal heterogeneity of extracellular cues—is critical for various physiological functions, including development, immune response, and wound healing (Ridley et al., 2003; Bear and Haugh, 2014; SenGupta et al., 2021). Ineffectual migration in response to external stimuli has been linked to developmental disorders (Guarnieri et al., 2018), immune deficiencies (Bouma et al., 2009), and chronic wounds (Wilkinson and Hardman, 2020). Aberrant migration, whether excessive or misguided, has been implicated in multiple diseases, most notably cancer metastasis (Bravo-Cordero et al., 2012). Chemotaxis, or cell migration biased by soluble chemical gradients, has been intensely studied over the last half-century, leading to important insights regarding the underlying signaling networks and cytoskeletal effectors that link receptor-mediated gradient sensing to directional movement. Initial studies in both fast-moving, ameboid-like cells (Servant et al., 2000) and slow-moving, mesenchymal cells (Haugh et al., 2000) showed that chemotactic gradients induce polarized phosphoinositide 3-kinase (PI3K) signaling, which is associated with local activation of the small GTPase Rac1. Mutual dynamic feedback regulates coincident PI3K/Rac1 signaling (Weiner et al., 2002; Johnson et al., 2015; Town and Weiner, 2023), promoting recruitment and activation of the WAVE and Arp2/3 complexes at the cells’ leading edges. Arp2/3-driven polymerization of branched actin arrays (Krause and Gautreau, 2014) drives membrane protrusion in migrating cells (Weiner et al., 1999).

Although receptor-mediated control of the PI3K/Rac1/Arp2/3 signaling module offers a satisfying explanation of chemotactic sensing and response, multiple studies have challenged this paradigm. At least in certain contexts, PI3K (Ferguson et al., 2007; Melvin et al., 2011), Rac1 (Monypenny et al., 2009), and the Arp2/3 complex (Wu et al., 2012; Leithner et al., 2016) have each proven to be dispensable for chemotaxis. Moreover, we identified a different pathway, with phospholipase C-γ1 (PLC-γ1) activation resulting in non-canonical phosphorylation of myosin light chain by protein kinase Cα (PKCα), as required for fibroblast chemotaxis to platelet-derived growth factor-BB (PDGF) (Asokan et al., 2014). Though clearly necessary for chemotactic navigation, it is not yet clear how and in what other contexts PLC-γ1 signaling biases asymmetric movement. Is PLC-γ1 signaling, like PI3K (Idevall-Hagren et al., 2012; Town and Weiner, 2023) and Rac1 (Wu et al., 2009; Johnson et al., 2015) activation, sufficient to initiate protrusion?

Whereas multiple tools exist to synthetically activate PI3K and Rac1 in live cells, none have yet been developed for PLC-γ1. PLC-γ1 is one of 13 mammalian PLC-family isozymes, which hydrolyze the plasma-membrane lipid substrate, phosphatidylinositol 4,5-bisphosphate (PIP_2_), to produce the lipid and cytosolic second messengers, 1,2-diacylglycerol (DAG) and inositol 1,4,5-trisphosphate (IP_3_), respectively (Gresset et al., 2012). The primary sequence of PLC-γ1 includes an X-Y linker region, which harbors multiple domains that participate in intra- and inter-molecular interactions; among these are regulatory Src homology 2 (SH2) domains that serve to autoinhibit enzymatic activity in unstimulated cells and mediate relief of autoinhibition in response to receptor stimulation (Hajicek et al., 2013, 2019) (**Fig. 1A**). PLC-γ1 is activated downstream of receptor tyrosine kinases, G protein-coupled receptors, the T-cell receptor, and β1 integrin (Jones et al., 2005; Tvorogov et al., 2005; Yang et al., 2012), and receptor activation initiates an activation program in which PLC-γ1 is recruited to the plasma membrane and phosphorylated on Tyr783. Tyr783 phosphorylation stabilizes a structural rearrangement of the enzyme that allows the catalytic core to interact with the membrane and hydrolyze PIP_2_. Unsurprising given its tightly coordinated regulation, both PLC-γ1 overexpression and activating mutations have been implicated in progression of cancer (Mandal et al., 2021) and other diseases (Tao et al., 2023). A deeper understanding of both lipase-dependent and-independent signaling mechanisms will be critical to contextualize and target the roles of PLC-γ1 in directed cell motility and in broader disease states (Hou et al., 2025).

**Figure 1:**
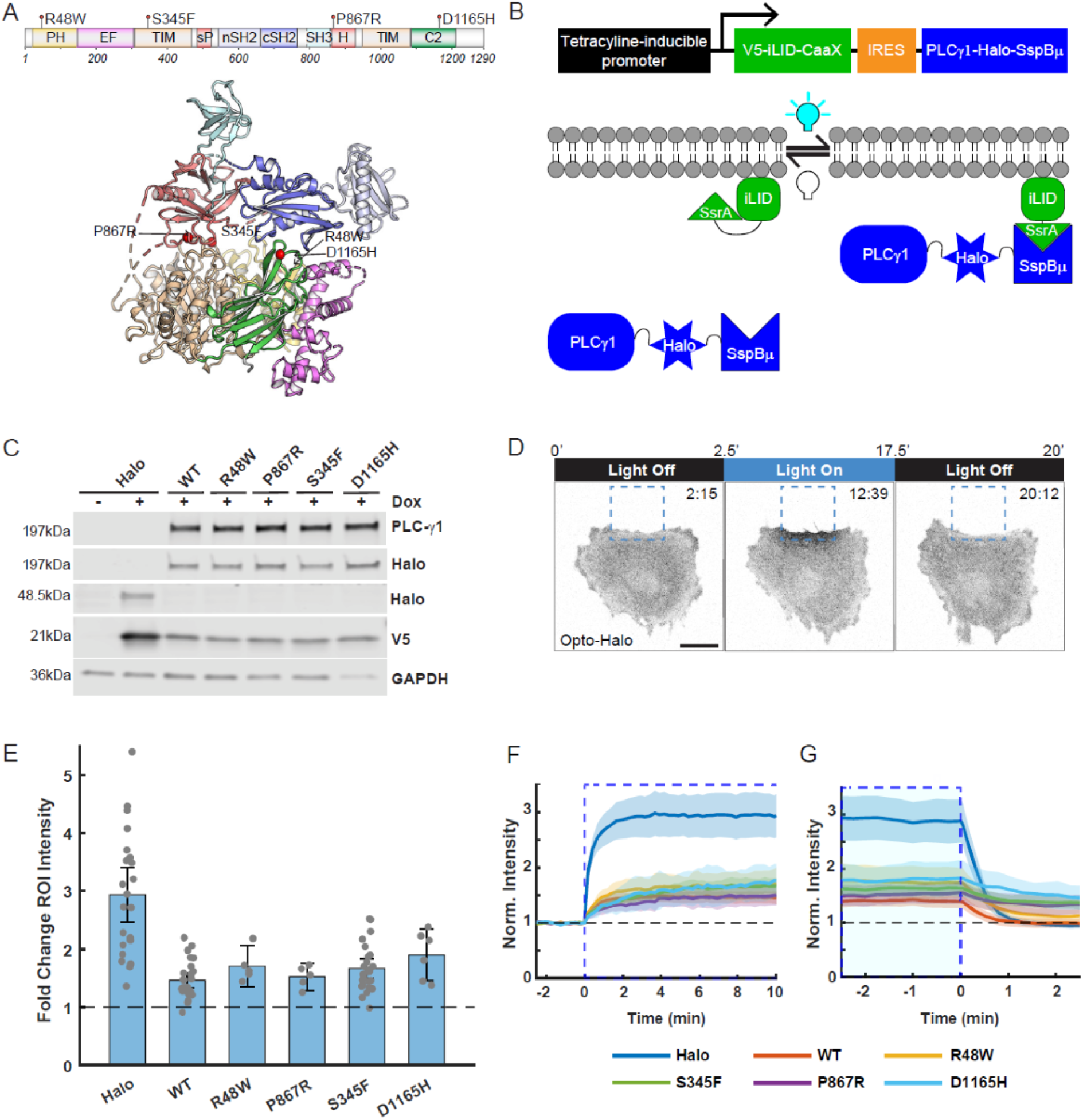
Inducible expression of OptoPLC-γ1 enables spatiotemporal control of membrane recruitment. **(A)** PLC-γ1 domain architecture and crystal structure showing the positions of activating mutations. **(B)** OptoPLC-γ1 expression is doxycycline-inducible, and blue light triggers membrane recruitment via the iLID-micro binding pair. **(C)** Immunoblot of V5-iLID-CaaX and Halo-SspBµ or PLCγ1-Halo-SspBµ variants inducibly expressed in *Plcg1*-null fibroblasts. The blot is representative of n = 3 biological replicates. **(D)** Time-lapse confocal micrographs of OptoHalo cells labeled with JF646 Halo ligand. The dashed, blue box marks the region of interest (ROI) illuminated with a 488-nm laser. Time indicates min:sec, and scale bar = 20 µm. Micrographs are representative of n = 22 cells. **(E)** Quantification of fold change of Halo-JF646 fluorescence intensity within the ROI upon photoactivation of OptoHalo or OptoPLC-γ1 variant. The fold change was calculated by fitting the fluorescence time-course to an exponential plateau function. Data are reported as the mean ± 95% confidence interval. **(F-G)** Time-course of Halo-JF646 enrichment within the photoactivated ROI in *Plcg1*-null fibroblasts rescued with the indicated OptoHalo/OptoPLC-γ1 variants upon focal illumination with a 488 nm laser. The time axes in (B) and (C) are scaled relative to the start and stop of focal illumination, respectively. Solid, colored lines represent the mean of the respective intensities for OptoHalo (n = 22 cells), OptoPLC-γ1 WT (n = 26 cells), OptoPLC-γ1 R48W (n = 5 cells), OptoPLC-γ1 P867R (n = 5 cells), OptoPLC-γ1 S345F (n = 25 cells), and OptoPLC-γ1 D1165H (n = 6 cells), and the shaded band regions indicate the 95% confidence interval.

Here, we describe the design and use of an optogenetic approach to control membrane recruitment of wild-type (WT) and gain-of-function PLC-γ1 variants in a *Plcg1*-null fibroblast-derived cell line. With this OptoPLC-γ1 system, we quantify the variants’ light-controlled membrane recruitment, regulatory phosphorylation, enzymatic activity, and stimulation of cell protrusion/motility polarization, ultimately showing that membrane recruitment of the cancer-associated mutant S345F induces lipase-dependent polarization of cell motility and persistent cell migration.

## RESULTS

### Construction of a light-controlled OptoPLC-γ1

To elucidate the role of PLC-γ1 in regulating cell motility, we sought to establish spatiotemporal control of enzymatic activity (**Figure 1**). Our engineered tool, OptoPLC-γ1, sandwiches a HaloTag (Encell et al., 2012) between full-length rat PLC-γ1 and the SspBµ component of the improved Light-Induced Dimer (iLID) (Guntas et al., 2015) system. Meanwhile, the light-sensitive iLID module is anchored to the plasma membrane via a CaaX domain. Upon blue light stimulation, the iLID module unfolds to expose SsrA, a peptide with high affinity for SspBµ (**Fig. 1B**). Wild-type (WT) PLC-γ1 is basally autoinhibited (Hajicek et al., 2013, 2019), and prior studies suggest that constitutive membrane targeting is insufficient for enzyme activation in cells (DeBell et al., 1999; Le Huray et al., 2022). We therefore elected to screen a panel of PLC-γ1 variants with weakly (R48W), moderately (P867R), or strongly (S345F, D1165) activating mutations (Hajicek et al., 2019) (**Fig. 1A**). To avoid cellular adaptation to constitutive, exogenous expression of mutant PLC-γ1, we constructed a single lentiviral vector enabling inducible, bicistronic expression of the OptoPLC-γ1 components. We established a panel of stable cell lines by infecting *Plcg1*-null fibroblasts (Ji et al., 1997) with the OptoPLC-γ1 variants, along with a Halo-SspBµ control (OptoHalo), and demonstrated that doxycycline treatment induces expression of both V5-iLID-CaaX and the Halo-SspBµ (**Fig. 1C**). Whereas V5-iLID-CaaX is expressed at a higher level in cells expressing OptoHalo relative to the OptoPLC-γ1 cell lines (attributable to use of the internal ribosomal entry site (IRES) to drive bicistronic expression (Hennecke et al., 2001)), expression levels of both proteins are consistent among the OptoPLC-γ1 lines (**Fig. 1C**) and localized as expected across all cell lines (**Supplemental Fig. S1A**).

To demonstrate spatiotemporal control of membrane recruitment, we imaged doxycycline-induced OptoPLC-γ1/OptoHalo cells labeled with JF646 HaloTag ligand on a laser-scanning confocal microscope; in each of these experiments, a peripheral region of interest (ROI) was focally illuminated using a 488-nm laser (**Fig. 1D**). Photoactivation of iLID yielded rapid and reversible enrichment of OptoPLC-γ1/OptoHalo (**Fig. 1E, F&G**). Among the cell lines, OptoHalo showed the greatest enrichment (consistent with its higher expression of V5-iLID-CaaX), whereas each of the OptoPLC-γ1 variants exhibited similar degrees of local, light-controlled recruitment.

### Acute membrane recruitment of PLC-γ1 with moderately/strongly activating mutation induces PIP_2_ hydrolysis and downstream signaling

Having established light-controlled recruitment of PLC-γ1, we sought to evaluate if membrane recruitment is sufficient to trigger PLC-γ1 activation. Given that phosphorylation on Tyr783 (pTyr783) is a hallmark of PLC-γ1 activation, we first evaluated basal and light-induced levels of pTyr783 in our OptoPLC-γ1 lines by immunoblotting (**Fig. 2A&B**). As a positive control and standard for comparison, we stimulated OptoPLC-γ1 WT cells with 1 nM PDGF, a known activator of PLC-γ1 signaling (Asokan et al., 2014). We observed elevated basal pTyr783 in cells expressing the highly active S345F and D1165H mutants, consistent with a recent study (Zeng et al., 2024). Cells expressing the moderately active P867R mutant also showed elevated basal pTyr783, whereas cells expressing the mildly active R48W mutant had levels comparable to WT. In response to blue light, pTyr783 levels in cells expressing OptoPLC-γ1 WT, P867R, S453F, or D1165H significantly increased, suggesting that membrane recruitment further relieves their autoinhibition of enzymatic activity; however, in WT cells the light-stimulated increase was much smaller than that stimulated by PDGF.

**Figure 2:**
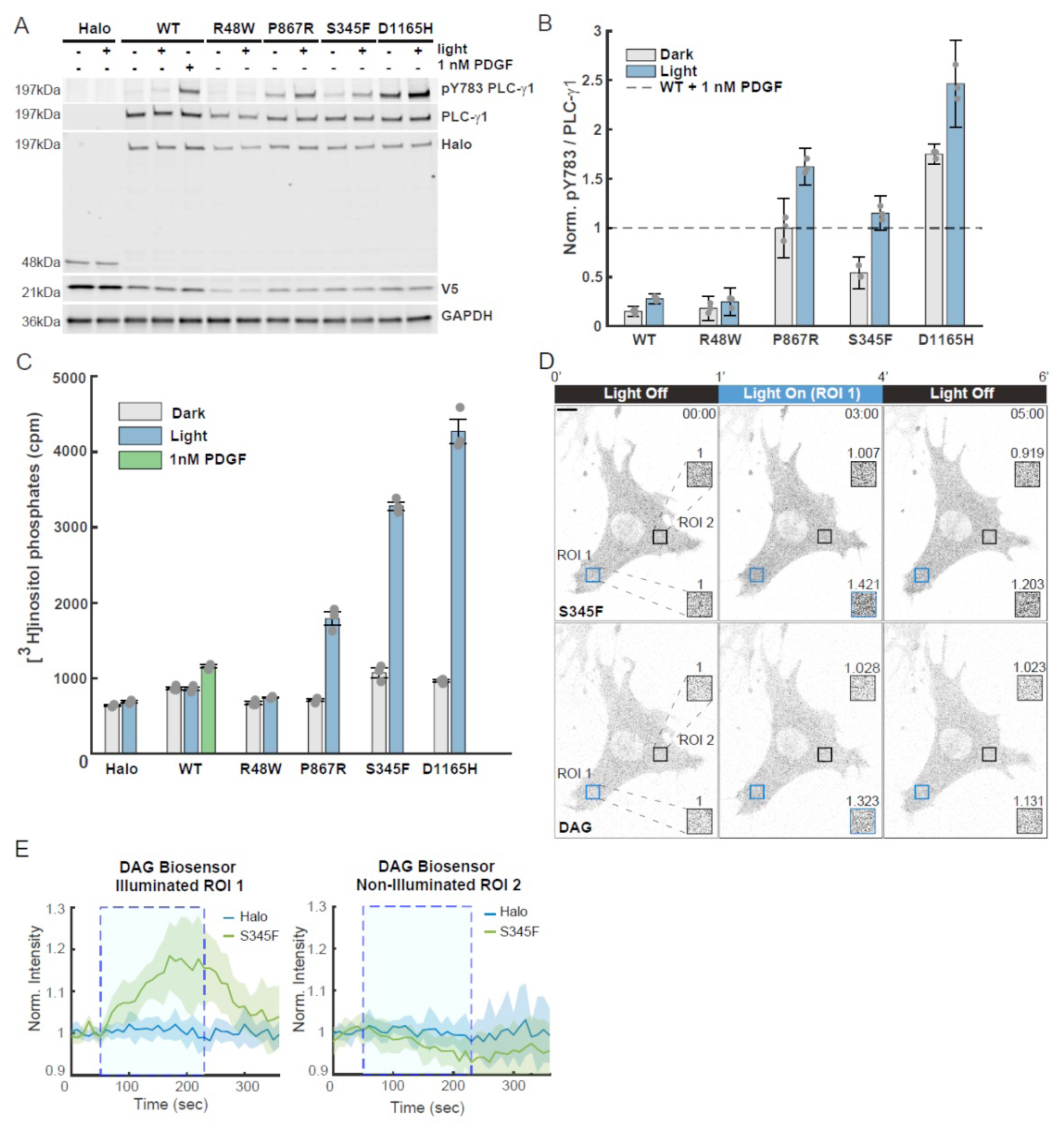
Membrane recruitment of OptoPLC-γ1 triggers enzyme activity. **(A)** Immunoblot of PLC-γ1 and associated pTyr783 in OptoHalo and OptoPLC-γ1 variant cells, globally stimulated with blue light or 1 nM PDGF for 5 minutes. The blot is representative of n = 3 biological replicates. **(B)** Quantification of pTyr783/PLC-γ1 ratio from immunoblots represented in **A**, normalized by that of PDGF-treated WT PLC-γ1. The data are reported as the mean ± 95% confidence interval. **(C)** Phospholipase activity (PIP_2_ hydrolysis) in OptoHalo/OptoPLC-γ1 cells globally stimulated with light or 1 nM PDGF for 15 minutes. The presented data are the mean ± SEM of triplicate samples from a representative experiment among n = 3 biological replicates. **(D)** Time-lapse confocal micrographs showing photoactivation of *Plcg1*-null fibroblasts expressing both OptoPLC-γ1 S345F and a DAG biosensor (PKCγ(C1A)-NES-mScarlet). Time indicates min:sec, and scale bar = 10 µm. Micrographs are representative of n = 7 cells **(E)** Time-course of DAG biosensor enrichment within the illuminated ROI (ROI 1, blue) and non-illuminated ROI (ROI 2, black) in *Plcg1*-null fibroblasts expressing the aforementioned DAG biosensor and rescued with either OptoHalo (n = 8 cells) or OptoPLC-γ1 S345F (n = 7 cells). Solid, colored lines represent the mean of the respective intensities, and the shaded band regions indicate the 95% confidence interval.

Another outcome of membrane localization is to position PLC-γ1 in proximity to its lipid substrate, PIP_2_. To assess the combined effect of membrane recruitment, we quantified OptoPLC-γ1-catalyzed PIP_2_ hydrolysis by measuring accumulation of inositol phosphates following lithium treatment (Waldo et al., 2010; Hajicek et al., 2019) (**Fig. 2C**). Here again, cells expressing OptoHalo and PDGF-treated cells expressing OptoPLC-γ1 WT are the negative and positive control, respectively. Global photoactivation of iLID in OptoPLC-γ1 WT and R48W cells did not appreciably increase PIP_2_ hydrolysis relative to unstimulated cells. In contrast, global photoactivation of iLID in the cells expressing OptoPLC-γ1 P867R, S345F, or D1165H stimulated PIP_2_ hydrolysis, to levels far exceeding that of OptoPLC-γ1 WT cells stimulated by PDGF. For our panel of cell lines, we corroborated the photoactivation of PLC-γ1 signaling at the level of Protein Kinase D1 (PKD) phosphorylation on Ser744/748, a known substrate of PKC isozymes (Waldron and Rozengurt, 2003; Kim et al., 2024) (**Supplemental Fig. S1B&C**). For OptoPLC-γ1 S345F, light stimulation of intracellular calcium rise is further corroboration of downstream signaling (**Supplemental Fig. S1D&E**). Interestingly, although pTyr783 and PIP_2_ hydrolysis readouts are qualitatively consistent, they are not well-correlated (compare **Fig. 2B&C**). In the absence of stimulation, cells expressing OptoPLC-γ1 P867R, S345F, or D1165H do not have substantially elevated basal PIP_2_ hydrolysis rates, despite possessing elevated pTyr783. Accordingly, light stimulation of iLID in each of those cell lines yields PIP_2_ hydrolysis that is far more striking than the corresponding increase in pTyr783. Additionally, cells expressing OptoPLC-γ1 S345F exhibited greater light-stimulated lipase activity, but lower light-stimulated pTyr783, than cells expressing OptoPLC-γ1 P867R. Together, these data support the concept that the combination of pTyr783 and membrane localization unlock the enzymatic potential of each PLC-γ1 variant. For the P867R, S345F, and D1165H mutants, light-controlled membrane recruitment is sufficient for PIP_2_ hydrolysis and downstream signaling; for WT and weakly activated R48W PLC-γ1, it is not.

To confirm that focal photoactivation of OptoPLC-γ1 enables spatiotemporal control of lipid hydrolysis, we measured DAG dynamics using a previously described translocation biosensor (Schuhmacher et al., 2020) in *Plcg1*-null fibroblasts expressing the highly active OptoPLC-γ1 S345F mutant. We then simultaneously measured the fluorescence intensities of OptoPLC-γ1 S345F and DAG biosensor in an illuminated ROI (ROI 1), using a non-illuminated ROI (ROI 2) as a control. While both OptoPLC-γ1 S345F and DAG fluorescence intensities increased in the illuminated ROI, indicating both PLC-γ1 recruitment and localized DAG accumulation, no changes in fluorescence intensities were observed in the non-illuminated ROI (**Fig. 2D&E and Supplemental Fig. S1F&G**). Experiments using *Plcg1-*null fibroblasts expressing both OptoHalo and the aforementioned DAG biosensor show an increase in Halo, but not DAG fluorescence intensity, in the illuminated ROI (**Fig. 2E** and **Supplemental Fig. 1F&G**) Together, these findings demonstrate that focal recruitment of OptoPLC-γ1 S345F results in localized conversion of PIP_2_ to DAG.

### Focal recruitment of PLC-γ1 with moderately/strongly activating mutation induces membrane ruffling and protrusion

Given previous reports implicating PLC-γ1 signaling as a key node in directed cell motility and cancer metastasis (Asokan et al., 2014; Sala et al., 2008), we asked if focal photoactivation of iLID in OptoPLC-γ1 cells could alter protrusion dynamics (**Figure 3**). Doxycycline-induced, JF646-labeled OptoPLC-γ1/OptoHalo cells were imaged and partially photoactivated as before, allowing space for sustained edge illumination should the cell protrude into the ROI (**Fig. 3A** and **Video 1**). To quantify the phenotypic responses to OptoPLC-γ1/OptoHalo photoactivation, we segmented photoactivated cells by manually thresholding the JF646 fluorescence intensity; by comparing the segmented cell masks prior to and at any time during photoactivation, net protruded/retracted areas are identified and measured (**Fig. 2B**). Reassuringly, focal blue-light illumination of OptoHalo control cells did not elicit appreciable net protrusion (**Supplemental Fig. S2A-C**). Among the OptoPLC-γ1 cell lines, neither WT nor R48W cells showed consistent responses. In contrast, light stimulation of cells harboring OptoPLC-γ1 P867R, S345F, or D1165H elicited visually striking protrusion within and proximal to the photoactivation ROI; and in those cells, regions opposite the photoactivated ROI often showed net retraction (**Fig. 3B**). To compare protrusion/retraction responses across the cohorts of photoactivated OptoPLC-γ1 cells, we track the time-dependent changes in area at the ‘front’ and ‘rear’ of photoactivated cells (5-6 per line). Setting the angular position 0° on the line between the ROI and cell centroids prior to photoactivation, we define ‘front’ and ‘rear’ regions as the 90° spans centered at 0° and 180°, respectively (**Fig. 3B&C**). The results show that light stimulation of the S345F mutant elicited the most robust cell polarization, with consistently large, positive and negative changes in area at the cell ‘front’ and ‘rear’, respectively (**Fig. 3C**).

**Figure 3:**
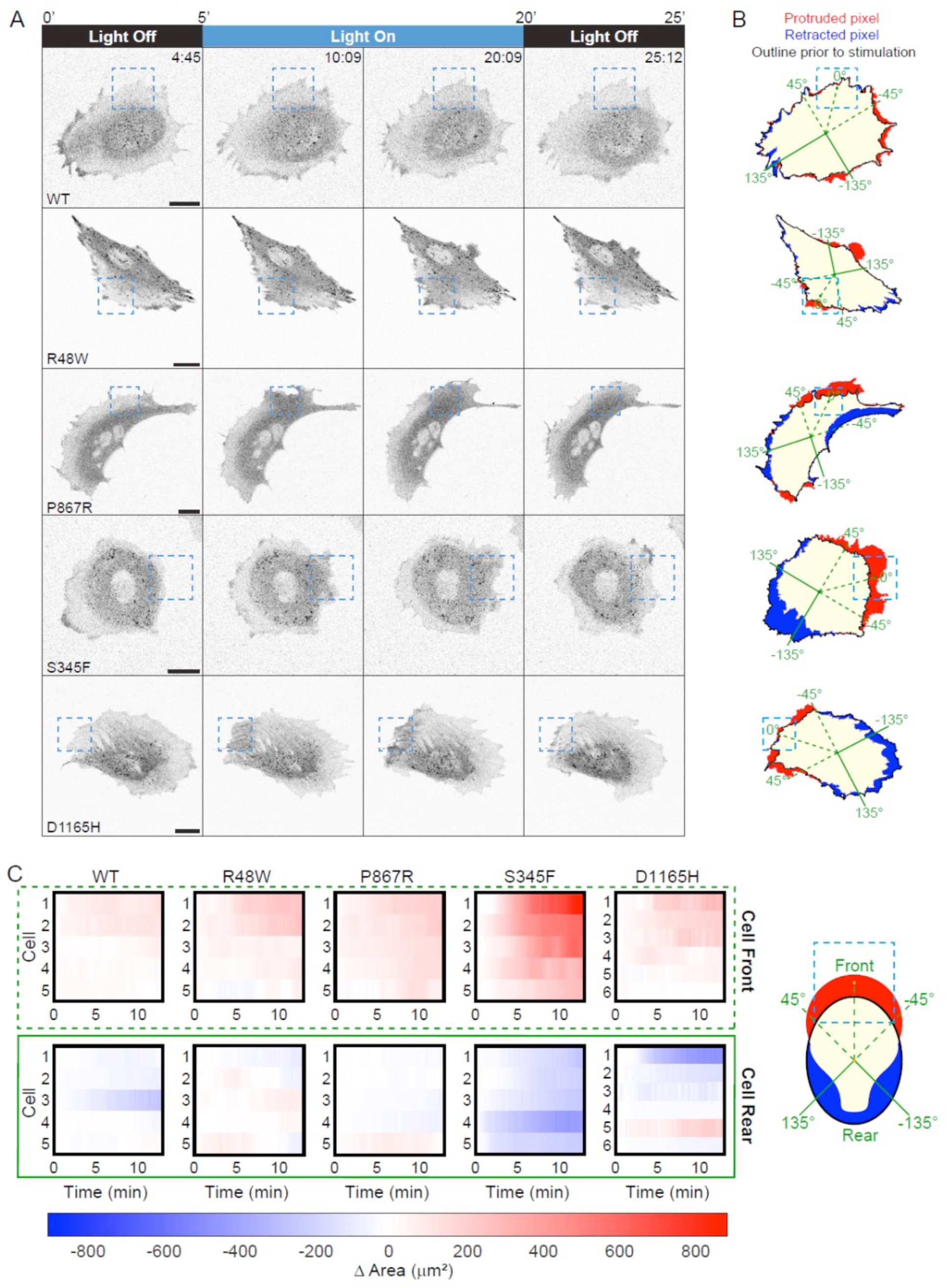
Focal photoactivation of active OptoPLC-γ1 mutants induces membrane ruffling and protrusion. **(A)** Time-lapse confocal micrographs of OptoPLC variant cells labeled with JF646 ligand and photoactivated as indicated. The dashed, blue box indicates the light-stimulated ROI. Time indicates min:sec, and scale bar = 20 µm. Selected cells are representative of n = 5 or 6 photoactivated cells for each cell line. **(B)** Pixel map highlighting protruded (red), retracted (blue), and common pixels (tan) between cell masks immediately prior to and the end of photoactivation for each variant shown in **A**. Green lines denote the angle positions for constructing spatiotemporal maps of edge velocity and binning changes in cell area. Solid black lines denote the outline of the cell mask immediately prior to photoactivation. **(C)** Heatmaps noting the change in area at the ‘cell front’ and ‘cell rear’ with respect to time for each photoactivated cell included in the initial screening cohort of OptoPLC-γ1 variant cell lines. The ‘cell front’ is defined as the 90-degree window centered on the line between the ROI centroid and cell centroid immediately prior to photoactivation as depicted by the cartoon (right). The ‘cell rear’ is defined as the 90-degree window mirroring the ‘cell front’. Time is scaled relative to the start of photoactivation.

An alternative visualization of a cell’s motility response is as a spatiotemporal protrusion/retraction map (Welf et al., 2012; Johnson et al., 2015), in which edge velocity as a function of angular position and time is estimated from the protruded/retracted pixels between frames in one-degree angular bins. Consistent with the visualizations of net protrusion/retraction described above, the corresponding velocity maps of OptoPLC-γ1 WT and R48W cells do not show appreciably altered edge dynamics; whereas those of OptoPLC-γ1 P867R, S345F, and D1165H cells show rapid changes, with waves of protrusion and retraction within and proximal to the photoactivated ROI (**Fig. S2D**). These waves are associated with visually observed membrane ruffling events (**Video 1**).

Spurred by these results, we asked if OptoPLC-γ1-induced protrusion occurs through mobilization of the Arp2/3-dependent, lamellipodial machinery. To that end, we expressed the highly active OptoPLC-γ1 S345F mutant in a previously described, tamoxifen-inducible *Arpc2* knockout (JR20) fibroblast line (Rotty et al., 2015; Chandra et al., 2022) and focally photoactivated control or tamoxifen-treated cells (**Fig. S2E-G).** As in *Plcg1-*null fibroblasts, focal photoactivation of OptoPLC-γ1 S345F in control JR20 fibroblasts induced protrusion, whereas this response was abolished in tamoxifen-treated cells. These data suggest that OptoPLC-γ1-induced lipid hydrolysis promotes protrusion through branched actin polymerization.

### OptoPLC-γ1 S345F photoactivation polarizes cell morphology and direction of motion on demand

Having identified OptoPLC-γ1 S345F as most responsive to local light stimulation, we asked if this mutant could induce polarization/repolarization of cell motility on demand, upon sequential or simultaneous photoactivation of distal regions (**Figure 4**). In one set of experiments, after photoactivating a cell expressing OptoPLC-γ1 S345F, we photoactivated the same cell again, with the ROI positioned elsewhere and the second light exposure initiated ∼20 minutes after cessation of the first. Strikingly, sequential recruitment of OptoPLC-γ1 S345 at different locations produced protrusion within and proximal to each ROI, while inducing retraction on the opposing side of the cell (**Fig. 4A&B, Supplemental Fig.S3A** and **Video 2**). The polarization and reversal of polarity occurred even when an ROI was placed on a site of inward curvature (**Fig. 4A, bottom**), which is considered refractory to protrusion (Welf et al., 2020; Sadhu et al., 2023; Town and Weiner, 2023), with an associated change in local morphology.

**Figure 4:**
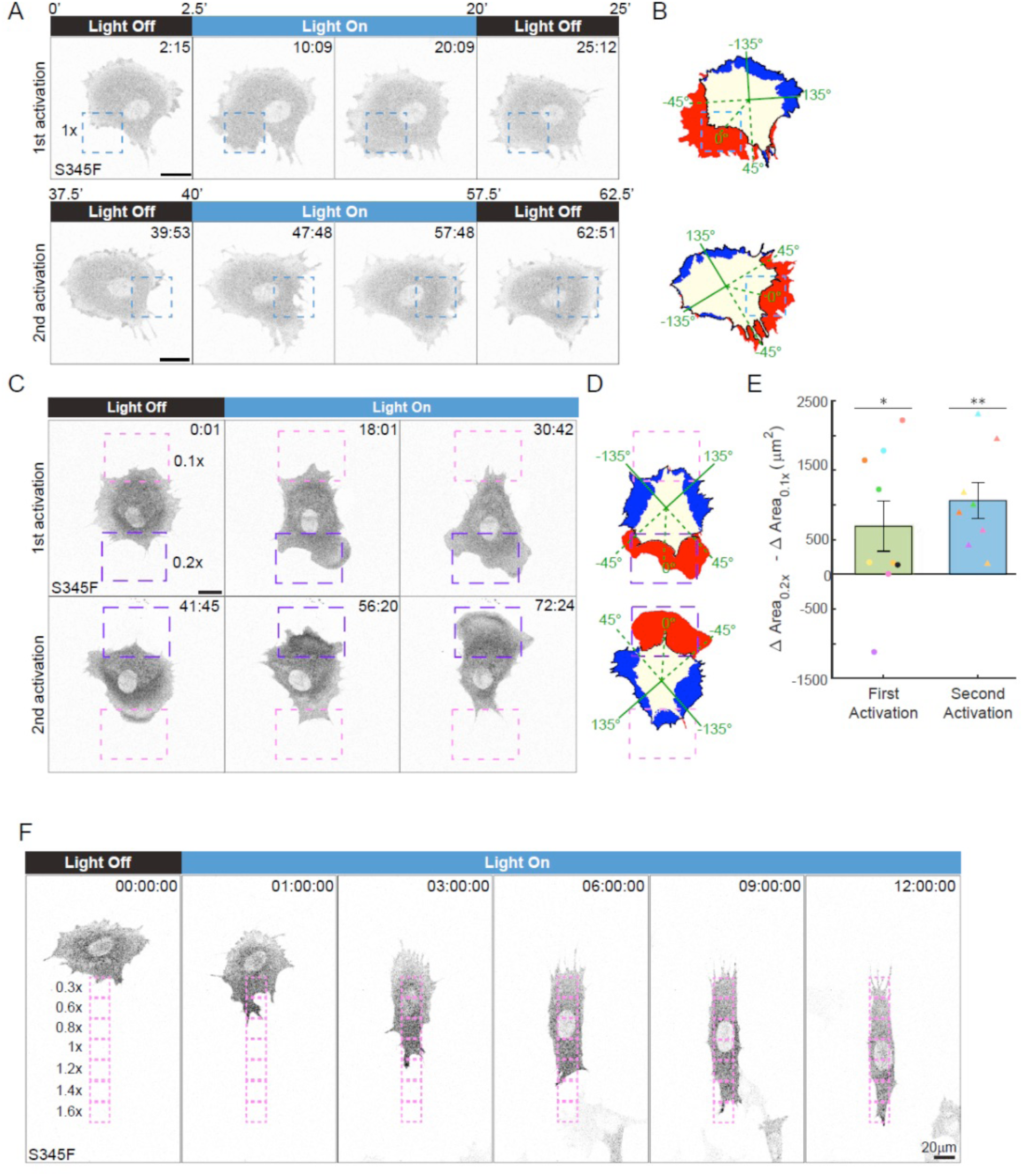
OptoPLC-γ1 S345F photoactivation repolarizes cell morphology and direction of motion. **(A)** Time-lapse confocal micrographs of an OptoPLC-γ1 S345F cell labeled with JF646 ligand, photoactivated as indicated. The same cell was subjected to a second photoactivation, with the illuminated ROI moved as indicated. Dashed, blue box indicates ROI illuminated with a 488-nm laser set to 0.1% (1x) power. Time indicates min:sec, and scale bar = 20 µm. Representative of n = 7 cells. **(B)** Net protrusion (red) and retraction (blue) for the first photoactivation (top) and second photoactivation (bottom) of the cell shown in **A** (presented as in Fig. 2B). **(C)**. Time-lapse confocal micrographs of an OptoPLC-γ1 S345F cell labeled with JFX554 ligand, photoactivated as indicated. The same cell was subjected to sequential photoactivation protocols with two different ROIs simultaneously illuminated with different laser powers, set at 0.02% (0.2x) and 0.01% (0.1x). The ROIs were moved between photoactivations to ensure that they still captured the cell edge, and the laser powers were flipped. Dashed, purple boxes indicate the 0.2x ROIs, and the dashed, pink boxes indicate the 0.1x ROIs. Time indicates min:sec, and scale bar = 20 µm. The sequence shown is representative of n = 8 cells. **(D)** Net protrusion (red) and retraction (blue) for the first (top) and second (bottom) photoactivations of the cell shown in **C**. **(E)** Quantification of differential change in area, calculated by subtracting the 0.1x change from the 0.2x change in area. Quantifications in (G-I) include 8 cells photoactivated twice sequentially (30-minute duration each, <10 minutes between) and a single cell, colored in black, photoactivated only once. Data are presented as the mean ± SEM. Delta area quantifications were compared against zero by the appropriate (right-tailed E) one-tailed t-tests. **: p < 0.01 and *: p < 0.05. **(F)** Time-lapse confocal micrograph of a cell expressing OptoPLC-γ1 S345F labeled with JF646 ligand and photoactivated as indicated. Time indicates h:min:sec, and scale bar = 20 µm. Representative of n = 6 cells.

We next asked if the photoactivation response of OptoPLC-γ1 S345F cells is sensitive to light dose. We first confirmed that local protrusions are still stimulated when the photoactivation laser power is reduced to 0.1x (**Supplemental Fig. S3B**). We then simultaneously illuminated two ROIs, situated on opposing sides of the cell, with 0.1x and 0.2x laser power. For a subset of responding cells, we subsequently repositioned the ROIs to maintain edge illumination and adjusted the ROI intensities to oppose the observed direction of motion (**Fig. 4C&D, Supplemental Fig.S3C** and **Video 3**). Analogous to the previous quantification of “front” and “rear” protrusion/retraction, we quantified cell polarization by measuring changes in area within and proximal to the 0.2x and 0.1x ROIs. Apart from one of the cells, the first photoactivation yielded positive changes in area within and proximal to the 0.2x ROI (**Supplemental Fig.S3D**). In contrast, on the 0.1x side, a mix of protrusion and retraction responses were observed (**Supplemental Fig S3E**). Interestingly, upon second photoactivation, all of the cells polarized directionality toward the 0.2x ROI, and the cells predominantly retracted on the 0.1x side (**Fig. 4E and Supplemental Fig S3D&E**).

Compelled by these results, we assessed whether or not OptoPLC-γ1 S345F photoactivation could elicit cell migration in a defined direction. *Plcg1-*null fibroblasts expressing OptoPLC-γ1 S345F were exposed to an optotactic gradient, presented as a series of adjacent ROIs, with stepwise increases in illumination intensity. Remarkably, OptoPLC-γ1 S345F-expressing cells consistently polarized motility and persistently migrated up the gradient (**Fig. 4F** and **Video 4**). Control experiments demonstrated that OptoHalo-expressing cells were unable to either promote protrusion or sustain migration when exposed to the same optotactic gradient (**Supplemental Fig. S3F** and **Video 5**). Altogether, these results demonstrate that dose-dependent, light-induced focal recruitment of OptoPLC-γ1 S345F robustly and reversibly polarizes cell morphology and directionality, enabling persistent cell migration.

### PKCα inhibition or calcium chelation attenuates, but does not block, the response to OptoPLC-γ1 S345F recruitment

Having characterized the OptoPLC-γ1 S345F photoactivation response, we sought to probe the underlying mechanism at the level of canonical pathways downstream of PIP_2_ hydrolysis (**Figures 5-7**). One is activation of PKC isozymes through increased abundance of the lipid product, DAG, in the plasma membrane(Black and Black, 2021). We previously demonstrated that the PLCγ1-DAG-PKCα pathway is required for mesenchymal chemotaxis (Asokan et al., 2014). We therefore hypothesized that PKC inhibition would block OptoPLC-γ1-driven protrusion (**Fig. 5A**). Prior to photoactivation, cells expressing OptoPLC-γ1 S345F were pretreated with either a vehicle control (DMSO) or 1 µM Gö6976, a conventional PKC inhibitor (Martiny-Baron et al., 1993) (**Fig. 5B&C** and **Video 6**). We confirmed that the inhibitor treatment is effective, as it blocks phorbol ester-stimulated phosphorylation of PKC substrates (**Supplemental Fig. S4A**). To our surprise, 1 µM Gö6976 did not abrogate membrane ruffling upon S345F photoactivation (**Fig. 5D**). Though reduced relative to the vehicle control, S345F photoactivation in 1 µM Gö6976-treated cells still elicited protrusion at the cell ‘front’, while retraction at the cell ‘rear’ was mostly blocked (**Fig. 5E&F**). To quantify the overall polarization response, we calculated the difference between the changes in area at the cell ‘front’ and ‘rear’. This metric further highlights that PKCα inhibition reduced, but did not block, S345F photoactivation-induced polarization (**Fig. 5G**).

**Figure 5:**
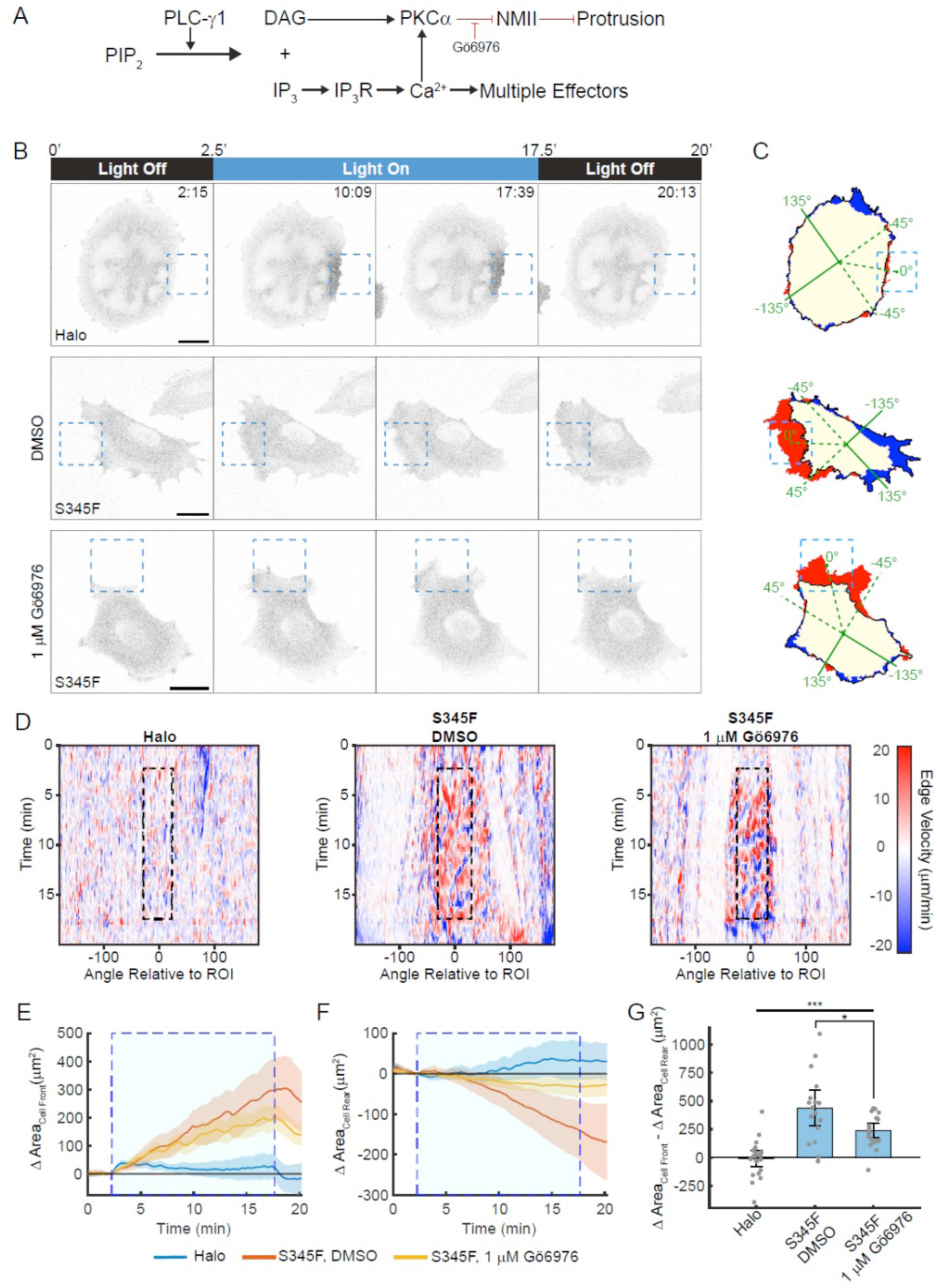
PKCα inhibition partially blocks the overall response of OptoPLC-γ1 S345F to light stimulation. **(A)** Schematic model of PLC-γ1 signaling and pharmacological inhibition of PKCα. **(B)** Time-lapse confocal micrographs of OptoHalo and OptoPLC-γ1 S345F cells, labeled with JF646 ligand and treated with either vehicle control (0.2% DMSO) or PKCα inhibitor, and photoactivated as indicated. Scale bars = 20 µm. Representative of n = 22 cells (Halo); n = 17 cells (S345F, DMSO); n = 20 cells (S345F, Gӧ6976). **(C)** Net protrusion (red) and retraction (blue) for the representative cells shown in **B**. **(D)** Spatiotemporal maps of edge velocity prior to, during, and after photoactivation for the cells shown in **B**. **(E&F)** Time-course of the change in area at the cell ‘front’ **(E)** and ‘rear’ **(F)**. The large, shaded blue rectangle denotes the period of photoactivation. Solid, colored lines represent the mean, and the shaded bands indicate the associated 95% confidence interval. **(G)** Quantification of polarized change in area, calculated by subtracting the ‘rear’ change in area from the ‘front’ change in area. Data are presented as the mean ± 95% confidence interval. Significance between groups assessed by ANOVA followed by Tukey-Kramer post-hoc test. ***: p < 0.001 and *: p < 0.05.

Another canonical PLC-γ1 signaling response is increased cytosolic calcium (Ca^2+^) (Clapham, 2007), which likewise has been implicated as a key regulator of directional motility (Wei et al., 2009; Tsai et al., 2014; Kim et al., 2016). We therefore asked if Ca^2+^ signaling is responsible for S345F-driven polarization (**Fig. 6A**). PLC-γ1-associated Ca^2+^ increase is achieved through both IP_3_ receptor (IP3R)-mediated release from intracellular stores and influx across the plasma membrane (Hofmann et al., 1999; Albert et al., 2008). Accordingly, in photoactivated OptoPLC-γ1 S345F cells, neither of two biochemically distinct IP3R antagonists (Gambardella et al., 2021) (10 µM xestospongin C (Gafni et al., 1997) or 100 µM 2-aminoethoxydiphenyl borate (2-APB) (Maruyama et al., 1997)) block the increase in cytoplasmic Ca^2+^, measured using the genetically encoded sensor, R-GECO (Kim et al., 2016); however, combining 2-APB treatment with EGTA (10 mM) to chelate extracellular Ca^2+^ does block the intracellular Ca^2+^ response, as does treatment with a cell-permeable Ca^2+^ chelator, BAPTA-AM (Tsien, 1981) (10 µM) (**Supplemental Fig. S4B&C**). Next, we photoactivated OptoPLC-γ1 S345F cells, pre-treated with either a vehicle control (DMSO) or 10 µM BAPTA-AM, and visualized protrusion dynamics as described above (**Fig. 6B-D** and **Video 7**). Quantification shows that, like PKCα inhibition, BAPTA-AM treatment reduced but did not block OptoPLC-γ1 S345F-driven protrusion at the cell ‘front’, while retraction at the cell ‘rear’ was mostly blocked (**Fig. 6E-G**).

**Figure 6:**
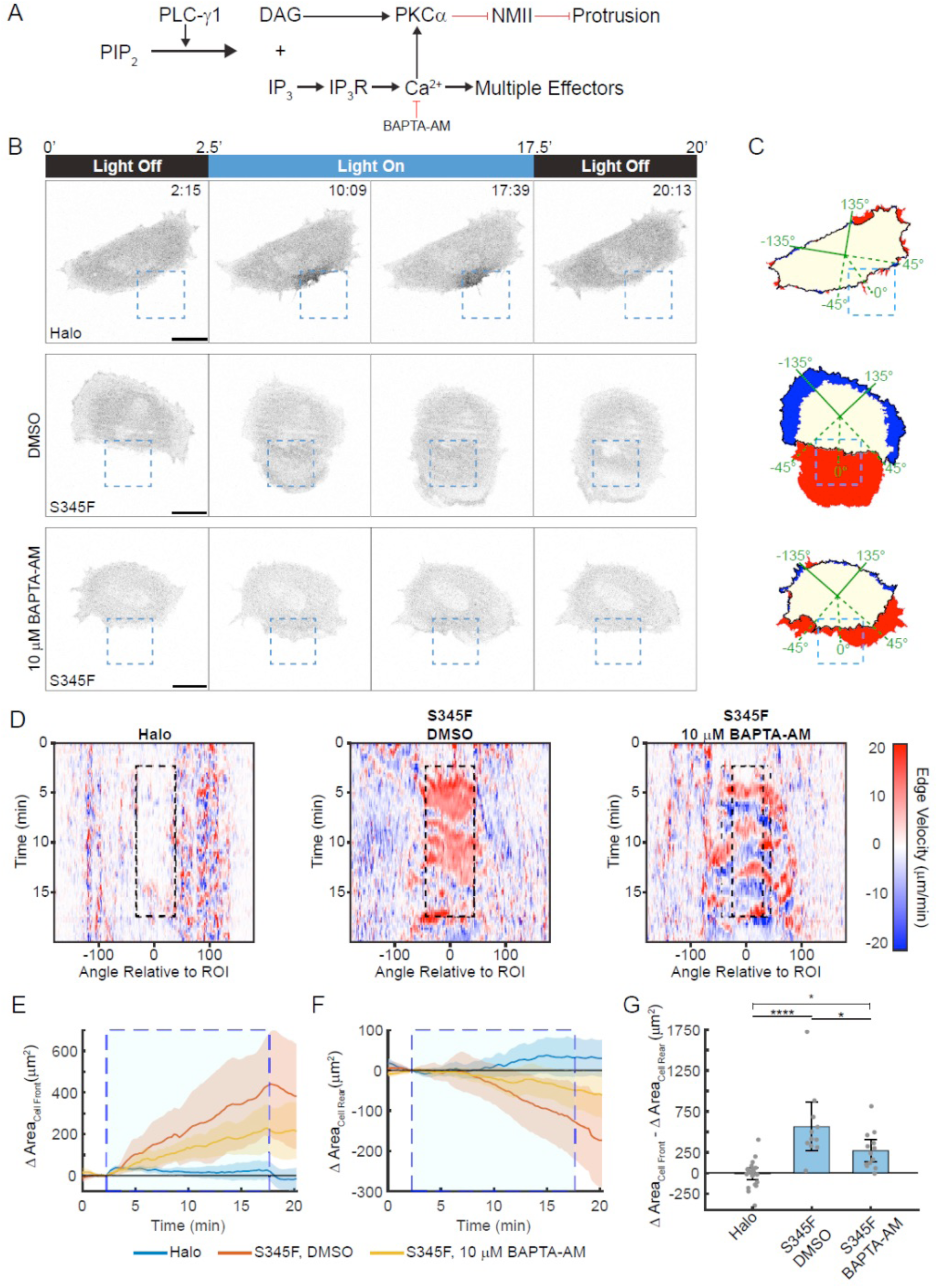
Calcium chelation hinders, but does not block, OptoPLC-γ1 S345F-induced repolarization. **(A)** Schematic model of PLC-γ1 signaling and pharmacological chelation of Ca^2+^. **(B)** Time-lapse confocal micrographs of OptoHalo or OptoPLC-γ1 S345F, labeled with JF646 ligand, photoactivated as indicated; OptoPLC-γ1 S345F were treated with either vehicle control (0.07% DMSO) or 10 µM BAPTA-AM. Scale bars = 20 µm. Representative of n = 22 cells (Halo); n = 11 cells (S345F, DMSO); n = 13 cells (S345F, BAPTA-AM). **(C)** Net protrusion (red) and retraction (blue) for the representative cells shown in **B**. **(D)** Spatiotemporal maps of edge velocity prior to, during, and after photoactivation for the associated cells shown in **B**. **(E&F)** Time-course analysis of change in area at the cell ‘front’ (**E**) and ‘rear’ (**F**). The large, shaded blue rectangle denotes the period of photoactivation. Solid, colored lines represent the mean, and the shaded bands indicate the associated 95% confidence interval. **(G)** Quantification of polarized change in area, calculated by subtracting the ‘rear’ change in area from the ‘front’ change in area. Data are presented as the mean ± 95% confidence interval. Significance between groups assessed by ANOVA followed by Tukey-Kramer post-hoc test. ****: p < 0.0001 and *: p < 0.05. For comparison, the OptoHalo data are the same as shown in Fig. 5E-G.

To evaluate the combinatorial effect of Ca^2+^ and PKC signaling on the protrusion response, we focally photoactivated cells expressing OptoPLC-γ1 S345F that had been pretreated with both 1 µM Gö6976 and 10 µM BAPTA-AM or a vehicle control. Similar to application of either inhibitor, photoactivation of cells expressing OptoPLC-γ1 S345F in the presence of the inhibitor combination induced protrusion at the cell front, albeit reduced relative to the vehicle control, while retraction at the cell ‘rear’ was mostly abolished (**Fig. 7**). We conclude that the two canonical ‘arms’ of PLC-γ1 signaling are dispensable for protrusion induced by local OptoPLC-γ1 S345F recruitment.

**Figure 7:**
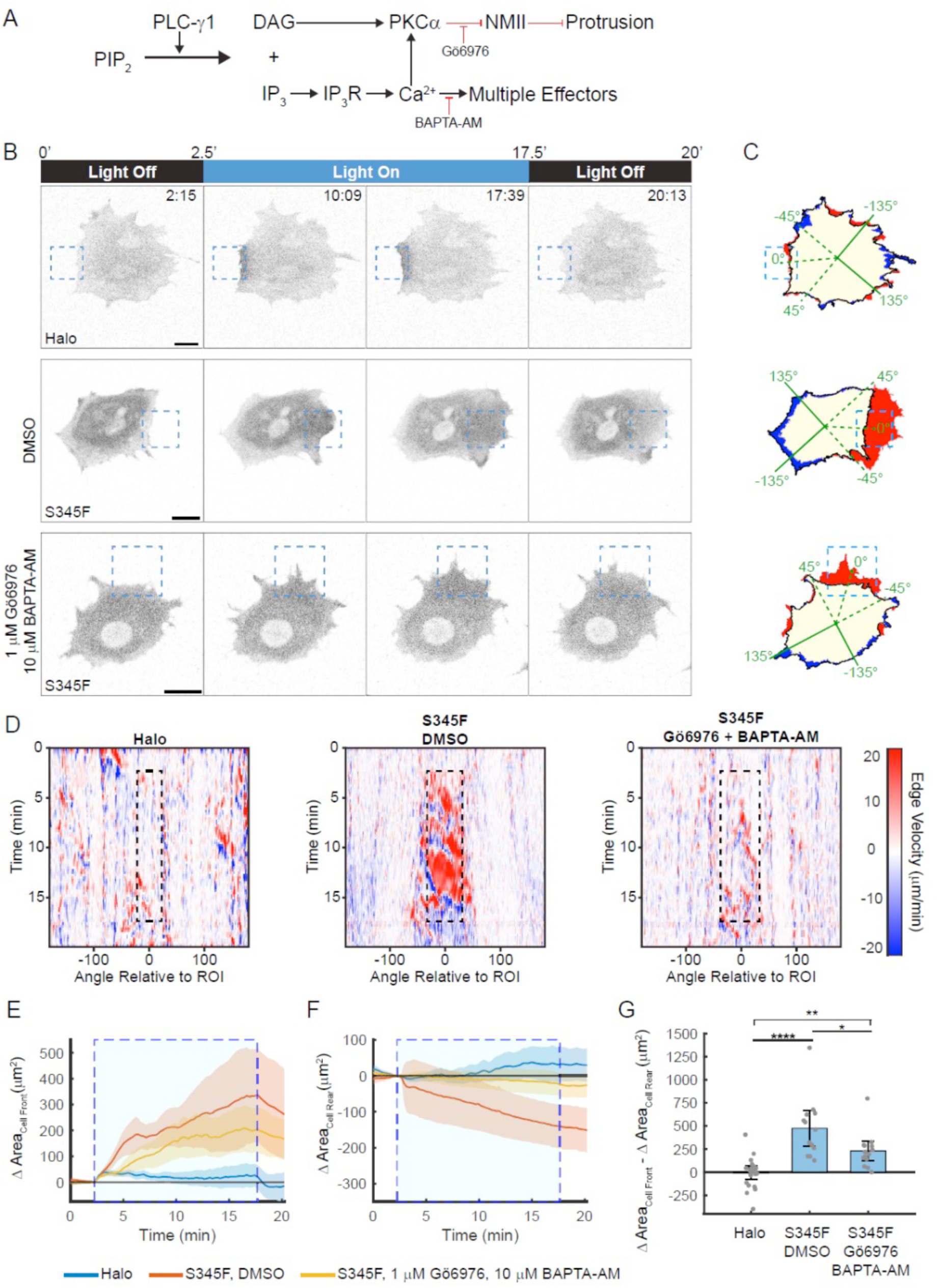
Combinatorial inhibition of PKCα and calcium signaling reduces, but does not block, protrusion upon OptoPLC-γ1 S345F photoactivation. **(A)** Schematic model of PLC-γ1 signaling and pharmacological inhibition of PKCα and chelation of Ca^2+^. **(B)** Time-lapse confocal micrographs of OptoHalo or OptoPLC-γ1 S345F, labeled with JF646 ligand, photoactivated as indicated; OptoPLC-γ1 S345F were treated with either vehicle control (0.1333% DMSO) or 1 µM Gӧ6976 and 10 µM BAPTA-AM. Scale bars = 20 µm. Representative of n = 22 cells (Halo); n = 13 cells (S345F, DMSO); n = 13 cells (S345F, Gӧ6976 & BAPTA-AM). **(C)** Net protrusion (red) and retraction (blue) for the representative cells shown in **B**. **(D)** Spatiotemporal maps of edge velocity prior to, during, and after photoactivation for the associated cells shown in **B**. **(E&F)** Time-course analysis of change in area at the cell ‘front’ (**E**) and ‘rear’ (**F**). The large, shaded blue rectangle denotes the period of photoactivation. Solid, colored lines represent the mean, and the shaded bands indicate the associated 95% confidence interval. **(G)** Quantification of polarized change in area, calculated by subtracting the ‘rear’ change in area from the ‘front’ change in area. Data are presented as the mean ± 95% confidence interval. Significance between groups assessed by ANOVA followed by Tukey-Kramer post-hoc test. ****: p < 0.0001 and *: p < 0.05. For comparison, the OptoHalo data are the same as shown in Fig. 5E-G.

### Impairing OptoPLC-γ1 S345F lipase activity impairs cell polarization

Having examined canonical signaling downstream of PIP_2_ hydrolysis, we next asked to what extent the observed photoactivation response depends on lipase activity (**Figures 8&9**). Hence, we sought to disable OptoPLC-γ1 S345F lipase activity by two parallel approaches (**Fig. 8A**). One was to render Tyr783 nonphosphorylatable via Y783F mutation, which essentially blocks WT PLC-γ1 activation (Kim et al., 1991) by precluding relief of autoinhibition (Gresset et al., 2010). As already shown, pTyr783 of OptoPLC-γ1 S345F is basally elevated and increases in response to light stimulation (Fig. 2B); however, the extent to which S345F mutation circumvents pTyr783 regulation of enzyme activity is unknown. Alternatively, the more direct approach is to target residues that directly affect catalysis. H335A is an established, lipase-dead PLC-γ1 mutant (Le Huray et al., 2022; Siraliev-Perez et al., 2022) that, based on structural analogy to PLC-δ, has been predicted to bind PIP_2_ but not hydrolyze it (Siraliev-Perez et al., 2022).

**Figure 8:**
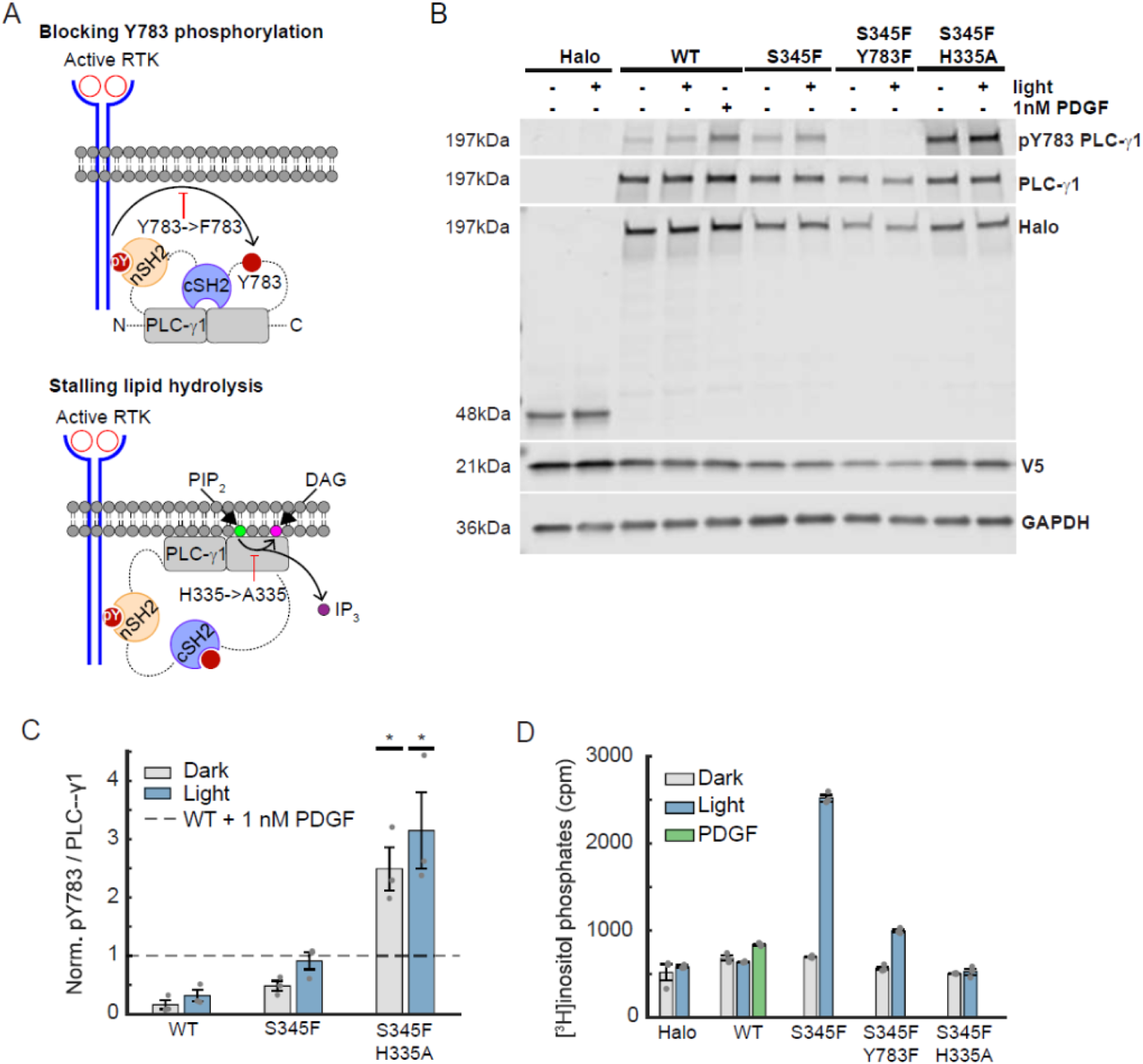
Blocking Tyr783 phosphorylation partially inhibits light-mediated OptoPLC-γ1 S345F lipase activity. **(A)** Schematic model illustrating the effects of Y783F and H335A single amino acid substitutions on canonical PLC-γ1 activation. **(B)** Immunoblot of PLC-γ1 and associated pTyr783 in OptoHalo and OptoPLC-γ1 variant cells, globally stimulated with blue light or 1 nM PDGF for 5 minutes. The blot is representative of n = 3 biological replicates. **(C)** Quantification of pTyr783/PLC-γ1 ratio from immunoblots represented in **B**, normalized by that of PDGF-treated WT PLC-γ1. The normalized data are reported as the mean ± SEM. The pTyr783 of S345F/H335A cells, both basal and light stimulated, were compared against the WT + 1 nM PDGF normalization by a one-tailed t-test. *: p < 0.05. **(D)** Phospholipase activity/PIP_2_ hydrolysis in OptoHalo/OptoPLC-γ1 cells globally stimulated with light, or PDGF, for 15 minutes. The presented data are the mean ± SEM of triplicate samples from a representative experiment among n = 3 biological replicates.

We characterized the S345/Y783F and S345F/H335A double mutants, relative to S345F and WT, in our OptoPLC-γ1 system. We confirmed doxycycline-induced, bicistronic expression of OptoPLC-γ1 in the corresponding cell lines (**Fig. 8B**), with localization comparable to OptoPLC-γ1 WT and S345F (**Supplemental Fig. S5A**), and we evaluated the effect of global photoactivation on pTyr783 (**Fig. 8B&C**) and PIP_2_ hydrolysis (**Fig. 8D**). The results confirm that pTyr is undetectable in cells expressing OptoPLC-γ1 S345F/Y783F, and we replicated the previous pTyr783 results for S345F relative to WT; however, to our surprise, we found that the S345F/H335A double-mutant is dramatically hyperphosphorylated, even basally (**Fig. 8D&E**). Treatment of the OptoPLC-γ1 lines with PDGF (2 nM) revealed that the basal pTyr783 levels are far from saturation, even in the S345F/H335A line (**Supplemental Fig. S5B**). With regard to enzymatic activity, the results confirmed that OptoPLC-γ1 S345F/H335A is lipase-dead; however, the cells expressing OptoPLC-γ1 S345F/Y783F exhibited reduced yet significant activation of PIP_2_ hydrolysis in response to light stimulation (**Fig. 8F**). Accordingly, assessment of downstream signaling showed that photoactivation of iLID in OptoPLC-γ1 S345F/Y783F cells, but not in OptoPLC-γ1 S345F/H335A cells, yields a small yet statistically significant increase in PKD phosphorylation on Ser744/748 (**Supplemental Fig. S5C&D**).

Having thus characterized these OptoPLC-γ1 variants, we transitioned to live-cell microscopy experiments and confirmed that focal photoactivation elicited reversible recruitment of both OptoPLC-γ1 S345F/Y783F and S345F/H335A, comparable to S345F (**Fig. 9A**). Interestingly, however, the reversible translocation responses among these cell lines (including OptoHalo control cells) exhibited different dissociation kinetics upon cessation of blue-light exposure (**Supplemental Fig. S5E&F**). Whereas OptoHalo translocation rapidly decays, as expected according to published properties of the iLID-micro interaction (Hallett et al., 2016; Zimmerman et al., 2016), the apparent lifetimes of OptoPLC-γ1 S345F variants’ membrane localization are longer and differ among them accordingly to lipase activity, not pTyr783. Consistent with these results, photoactivating cells expressing OptoPLC-γ1 S345F/Y783F elicited visible protrusion within and proximal to the ROI, whereas photoactivating OptoPLC-γ1 S345F/H335A cells yielded none (**Fig. 9A-C** and **Video 8**). Quantifying the change in area at the cell ‘front’ for the cohort of photoactivated cells reveals that photoactivating OptoPLC-γ1 S345F/Y783F produced protrusion, albeit reduced relative to the S345F mutant (**Fig. 9D**). Conversely, photoactivating cells expressing OptoPLC-γ1 S345F/H335A did not produce appreciable protrusion at the cell ‘front’, comparable to photoactivated WT and OptoHalo. Consistent with the cell ‘front’ observations, photoactivated OptoPLC-γ1 S345F/Y783 produced retraction at the cell ‘rear’, though significantly reduced relative to photoactivated S345F (**Fig. 9E**). Quantifying the difference between the change in area at the cell ‘front’ and ‘rear’ shows that photoactivating OptoPLC-γ1 S345F/Y783F weakly but significantly polarized those cells (**Fig. 9F**); by the same quantitative metrics, OptoPLC-γ1 S345F/H335A cells exhibited no discernable motility response to iLID photoactivation. These results, together with the previous characterization of other activating mutations of OptoPLC-γ1, indicate that the ability to photostimulate enzymatic activity is both required for, and to a large degree predictive of, cell polarization.

**Figure 9:**
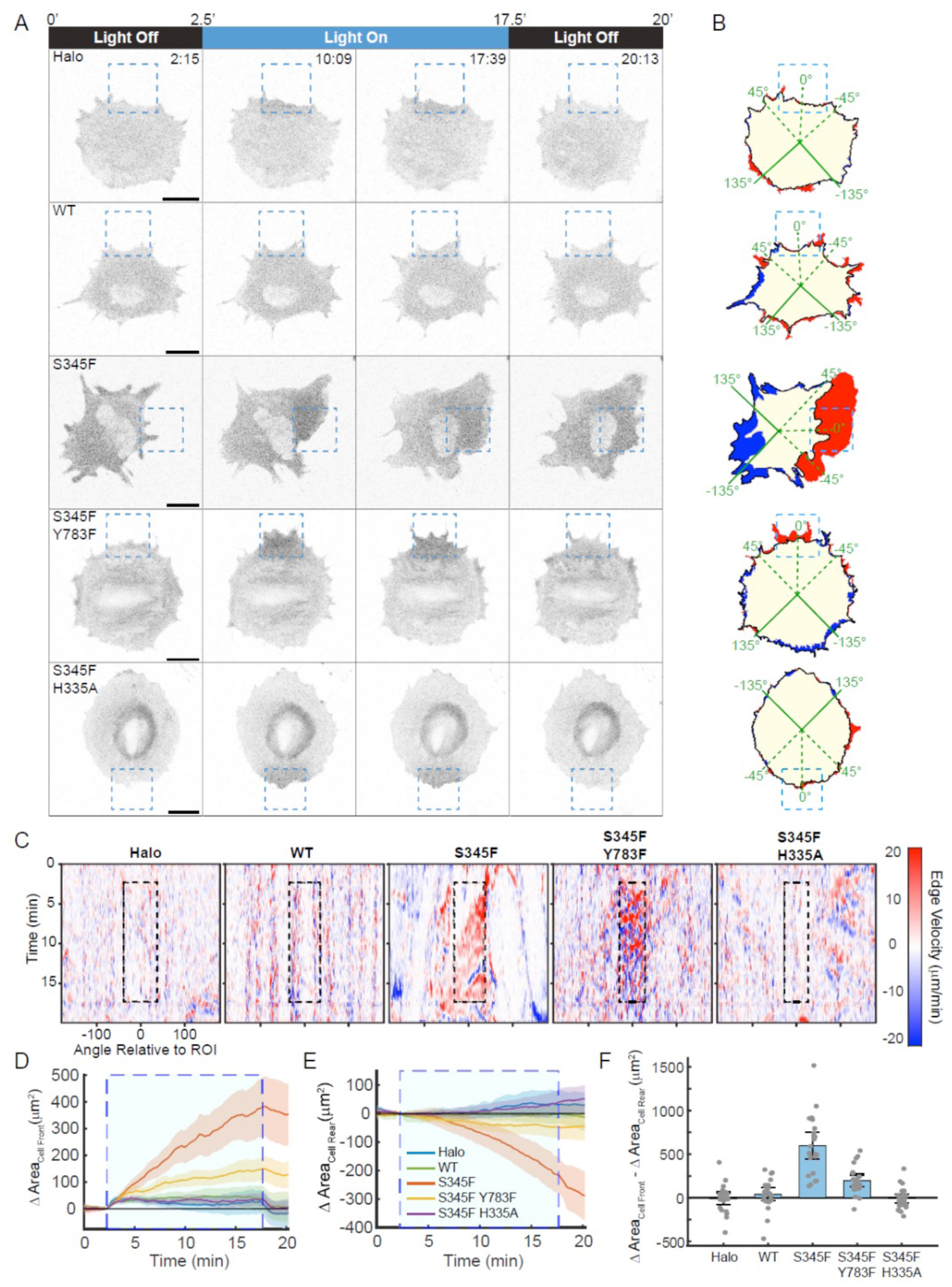
Cell polarization induced by OptoPLC-γ1 S345F is lipase-dependent. **(A)** Time-lapse confocal micrographs of OptoHalo and OptoPLC-γ1 variant cells, labeled with JF646 ligand and photoactivated as indicated. Scale bars = 20 µm. Representative of n = 22 cells (Halo), n = 23 cells (WT), n = 22 cells (S345F), n = 23 cells (S345F/Y783F), and n =24 cells (S345F/H335A). **(B)** Net protrusion (red) and retraction (blue), for the associated cells shown in **A**. **(C)** Spatiotemporal maps of edge velocity prior to, during, and after photoactivation for the associated cells shown in **A&B**. **(D&E)** Time-course analysis of change in area at the cell ‘front’ (**E**) and ‘rear’ (**F**). The large, shaded blue rectangle denotes the period of photoactivation. Solid, colored lines represent the mean, and the shaded bands indicate the associated 95% confidence interval. **(F)** Quantification of polarized change in area, calculated by subtracting the ‘rear’ change in area from the ‘front’ change in area. Data are presented as the mean ± 95% confidence interval. For comparison, the OptoHalo data are the same as shown in Fig. 5E-G.

## DISCUSSION

Seeking to dissect the role of PLC-γ1 in directing cell motility and to unveil aspects of PLC-γ1 enzymology in cells, we developed the OptoPLC-γ1 system. We successfully integrated cancer-associated OptoPLC-γ1 variants (Kataoka et al., 2015) to create many cell lines in a common PLC-γ1-null background, and we showed that light stimulation of lines expressing gain-of-function mutant OptoPLC-γ1s triggers PIP_2_ hydrolysis. In the same cell lines, photoactivation of a peripheral region of the cell rapidly elicited a local protrusion response. Moreover, we discovered that focal photoactivation of the OptoPLC-γ1 S345F mutant strikingly polarizes cells, with local protrusion accompanied by retraction on the opposite side of the cell, and that these cells can distinguish between regions of different intensities; when illuminated regions are arranged in a staircase gradient, the cells migrate persistently in the direction of increasing light intensity. So robust is the response of cells expressing OptoPLC-γ1 S345F to photoactivation, that blocking either or both of the two canonical PLC-γ1 signaling pathways or rendering Tyr783 nonphosphorylatable (S345F/Y783F) only partially attenuated it. Only upon eliminating the lamellipodial F-actin machinery, or rendering the mutant lipase-dead (S345F/H335A), was the motility response eliminated. Our results demonstrate the utility of the OptoPLC-γ1 system for dissecting PLC-γ1 regulation and function in cells, and its single-plasmid, inducible expression design may be adapted for recruitment of other proteins of interest. The addition of a HaloTag affords flexibility with respect to fluorescent labeling, which is particularly useful in optogenetic applications where the optical switch limits the available visible spectrum.

At the level of PLC-γ1 regulation and its perturbation by point mutations, we found satisfying corroboration of previous results (Hajicek et al., 2019) and new insights as well. For example, it was previously shown that the elevated lipase activity of PLC-γ1 R48W, relative to WT, is only apparent under strong receptor stimulation, suggesting that its gain-of-function attributes take effect after the enzyme is already active (Hajicek et al., 2019). Our light-stimulated recruitment data provide further insight into this notion, revealing that membrane proximity alone is insufficient to activate either OptoPLC-γ1 WT or R48W. Unlike the other mutations tested here, R48W is not located at the interface between the regulatory array and the catalytic core (Fig. 1A) and, perhaps due to its location, does not affect basally elevated Tyr783 phosphorylation. While Tyr783 phosphorylation is canonically viewed as a requirement for WT activation, it could also serve as a readout for disrupted autoinhibition, especially if the cSH2-pTyr783 interaction precludes pTyr783 dephosphorylation (Rotin et al., 1992). Notably, Tyr783 phosphorylation does not strictly correlate with light-mediated lipase activity; cells expressing OptoPLC-γ1 S345F exhibit greater light-mediated lipase activity than P867R, despite lower Tyr783 phosphorylation, and the lipase-dead S345F/H335A double mutant is hyperphosphorylated. Furthermore, rendering OptoPLC-γ1 S345F Tyr783-nonphosphorylatable reduces but does not block light-stimulated lipase activity. Taken together, these data are consistent with the concept that disruption of autoinhibition represents only one class of activating mutations, enhancement of membrane-binding affinity being another (Hajicek et al., 2019). Moving forward, the OptoPLC-γ1 system may be used to further test this concept, in conjunction with quantitative predictions generated from a kinetic model of PLC-γ1 activation (Nosbisch et al., 2022).

PLC-γ1 signaling has been implicated in cell spreading and migration (Tvorogov et al., 2005; Jones and Katan, 2007), cancer metastasis (Lattanzio et al., 2013; Jang et al., 2018), angiogenesis (Chen and Simons, 2021), and T-cell activation (Braiman et al., 2006)—processes in which multiple signaling pathways are integrated to modulate cytoskeletal dynamics. In the context of mesenchymal cell migration, there is a conundrum in that regard: although the PI3K/Rac1/Arp2/3 axis is sufficient to bias cell movement and is polarized in migrating fibroblasts, it is also dispensable for chemotactic responses of those cells. By contrast, PLC-γ1 signaling is both polarized and required for mesenchymal chemotaxis, but it was heretofore unclear if PLC-γ1 signaling is sufficient to bias cell movement. Here, we show that focal activation of PLC-γ1 is sufficient to induce localized membrane ruffling and protrusion. Furthermore, local photoactivation of the S345F mutant repolarizes cell morphology and redirects motion. Importantly, this repolarization is lipase-dependent, and the degree of dynamic motility correlates well with lipase activity induced by light, highlighted by our finding that cells expressing OptoPLC-γ1 S345F/Y783F double mutant exhibit partially reduced lipase activation and motility responses relative to OptoPLC-γ1 S345F. The response is also perhaps surprisingly robust to interventions that impair canonical PLC-γ1 S345F signaling. Blocking conventional PKCs, a treatment that prevents fibroblast chemotaxis to PDGF (Asokan et al., 2014), reduces but does not prevent the light-stimulated motility response; the same is true for even more extreme interventions, reducing intracellular calcium to an undetectable level or combinatorially blocking conventional PKC and Ca^2+^. These signaling aspects clearly contribute to, but are not absolutely required for, the light-induced motility response. What, then, are its other signaling determinants, and how are they integrated?

Among the many possible PLC-γ1 signaling mechanisms, including various non-canonical ones (Jones and Katan, 2007; Cooke and Kazanietz, 2022), the most direct is the local reduction of PIP_2_ abundance (Senju et al., 2017). Several studies support this notion. First, the concentration of free PIP_2_ in the membrane has been positively correlated with membrane tension/rigidity in fibroblasts (Raucher et al., 2000), which may be attributed to PIP_2_ interactions with ezrin/radixin/moesin (ERM) proteins anchoring the membrane to the actin cortex (Hao et al., 2009). Building on that concept, two recent studies have further implicated local reduction of membrane-cortex adhesion as permissive for initiation of membrane protrusion (Bisaria et al., 2020; Welf et al., 2020), and another line of research has linked membrane tension to the ability of amoeboid cells to polarize front and rear motility (De Belly and Weiner, 2024). Another longstanding motility mechanism attributed to PIP_2_ control is its ability to sequester cofilin (Zhao et al., 2010); release of cofilin in response to PLC-γ1 hydrolysis of PIP_2_ is implicated in local F-actin turnover and membrane protrusion in breast carcinoma cells (van Rheenen et al., 2007). Also important to consider is that PLC-γ1-dependent signaling does not function in isolation in cells. Once protrusion is initiated, so too are formation of adhesion complexes that mediate activation of other pathways. We envision that decoding mesenchymal motility will require a comprehensive understanding of PLC-γ1 signaling to the cytoskeleton and its integration with other signaling pathways, such as those affecting Arp2/3-mediated actin polymerization and NMII-dependent contractility. OptoPLC-γ1 can be instrumental in elucidating that integration.

## MATERIALS AND METHODS

### Constructs

All constructs were assembled according to standard multi-component Gibson Assembly protocols, utilizing the NEB HiFi Builder (New England Biolabs, Ipswich, MA) reagent. The pTLCV2-OptoPLCG1 construct was assembled as follows. The pTLCV2 plasmid was purchased from Addgene (Watertown, MA, plasmid # 87360 (Barger et al., 2019)) along with pCDNA5/FRT/HaloTag7-Geminin(1/110)_P2A_HaloTag9-Cdt(1-110)Cy- (plasmid # 169338 (Frei et al., 2022)). The U6 promoter and subsequent CRISPR sgRNA sequence was removed from the pTLCV2 parent plasmid, and the Cas9-2A-EGFP sequence was replaced by V5-iLID-CaaX-IRES-PLCG1-Halo-SspB(micro). The V5 sequence was appended via primer design to iLID-CaaX amplified from pLL7.0: Venus-iLID-CAAX (Guntas et al., 2015); the IRES sequence was amplified from pBM-puro-EGFP-AktPH (Melvin et al., 2011); full-length rat (*R. norvegicus*) *Plcg1* was amplified from pcDNA3.1-HA-PLCG1 (Hajicek et al., 2019); HaloTag7 was amplified from pCDNA5/FRT/HaloTag7-Geminin(1/110)_P2A_HaloTag9-Cdt(1-110)Cy-; and moderate-affinity SspB (micro) was amplified from pLL7.0: tgRFPt-SSPB R73Q (Guntas et al., 2015). pTLCV2-OptoPLCG1 variants were generated via site-directed mutagenesis. The pLL5.0-Lyn-R-GECO construct was assembled as follows. The p-Lyn-R-GECO (Kim et al., 2016) plasmid was purchased from Addgene (plasmid #120410) and inserted into the pLL5.0 backbone (Cai et al., 2007). The DAG biosensor, originally consisting of the C1A domain of PKCγ appended with a nuclear export signal and fused to EGFP (Schuhmacher et al., 2020), was a kind gift from Dr. André Nadler (Max Planck Institute of Molecular Cell Biology and Genetics); the construct was modified by replacing the EGFP tag with mScarlet.

### Inhibitors and growth factors

PDGF-BB (10014B10UG) was purchased from Peprotech (Cranbury, NJ). BAPTA-AM (2787/25), 2-APB (1224/10), and Gӧ6976 (2253) were purchased from Tocris (Minneapolis, MN). Xestospongin C (ab120914) was purchased from Abcam.

### Tissue culture, cell line generation and transfections

*Plcg1* (-/-) mouse embryonic fibroblasts (Ji et al., 1997) were a kind gift from Graham Carpenter. Cells were grown in high-glucose DMEM (4.5 g/L D-glucose, 0.584 g/L L-glutamate, 110 mg/L sodium pyruvate; Invitrogen) supplemented with 10% (v/v) fetal bovine serum (FBS, FB-01, Omega Scientific, Tarzana, CA), 1% non-essential amino acids (Invitrogen), and 1% glutamax (Invitrogen), in a 37 °C humidified incubator with 5% CO_2_. Stable cell lines were generated via lentiviral transduction. Briefly, lentiviral expression constructs were transiently co-transfected with pMDL-G/P-RRE (gag-pol), pRSV-REV, and pCMV-VSVG packaging plasmids into HEK293FT cells with X-tremeGene HP DNA transfection reagent (Roche/Sigma Millipore). Lentiviral supernatant was used to infect recipient cell lines in the presence of polybrene (sc-255611, Santa Cruz Biotechnology, Dallas, TX) at 3-5 µg/mL. To generate OptoHalo/OptoPLC-γ1 cell lines, infected *Plcg1-*null or JR20 fibroblasts were selected with 3 µg/mL or 2 µg/mL puromycin (A1113803, Thermo Fisher Scientific, Waltham, MA), respectively, then enriched for OptoHalo/OptoPLC-γ1 expression via fluorescence-activated cell sorting (FACS). Prior to sorting, cells were treated with doxycycline (D9891, Sigma-Aldrich, St. Louis, MO) (500 ng/mL for *Plcg1-*null or 1 µg/mL for JR20 fibroblasts) for 24 hours and labeled with 200 nM Janelia Fluor® JFX554 HaloTag® Ligand (HT1030, Promega, Madison, WI) per the manufacturer’s instructions. Following enrichment of cells exhibiting induced expression, *Plcg1-*null fibroblasts were subjected to an additional round of FACS to eliminate cells that lost response to doxycycline. Cells were maintained in puromycin (same as selection dose), and where appropriate, sequential rounds of FACS were performed to maintain cell lines expressing comparable levels of the desired constructs. Tamoxifen-induced *Arpc2* knockout in JR20 fibroblasts was performed as previously described. Briefly, JR20 fibroblasts harboring, but not expressing, OptoPLC-γ1 S345 were treated with 2 μM 4-hydroxy-tamoxifen (4-OHT). After two days, the media was replaced with a second dose of 4-OHT. Two days after the second dose, 4-OHT media was removed and cells were cultured in full media *sans* 4-OHT. Tamoxifen-treated cells were used in live-cell microscopy experiments between 7-10 days following the first dose of 4-OHT.

### Immunoblotting

Global photoactivation of OptoPLC-γ1 variants/OptoHalo was performed as follows: near-confluent dishes were positioned 12 inches away from an array of blue LED lights attached to the bottom side of a shelf in a CO_2_ incubator, maintained at 37°C. Cells were exposed to the blue light for 5 minutes and harvested immediately thereafter for immunoblotting analysis. With the exception of PKC inhibitor validation experiments, cells were lysed on ice in RIPA buffer containing 50 mM Tris, pH 8, 150 mM NaCl, 0.5% deoxycholate, 0.1% SDS, 1% NP-40, supplemented with protease and phosphatase inhibitor cocktails (87786 and 78420, respectively, Thermo Fisher, Scientific, Waltham, MA). For PKC inhibitor validation, cells were lysed on ice in 2x Laemmli sample buffer and sonicated for 10 seconds. Detection of phosphorylation and protein expression levels by immunoblotting was performed using the following antibodies: phospho-PLCG1(Y783) (14008, Cell Signaling Technology, Danvers, MA), PLCG1 (AB302940, Abcam, Cambridge, UK), Halo (G9211, Promega, Madison, WI), GAPDH (AM4300, Thermo Fisher Scientific, Waltham, MA), phospho-PKD (Ser744/748) (2054, Cell Signaling Technology, Danvers, MA), PKD (PA5-13749 Thermo Fisher Scientific, Waltham, MA), phospho-(Ser) PKC Substrate (2261, Cell Signaling Technology), phosphor-MLC(S1) (MP3461, ECM Biosciences, Davis, CA) and secondary antibodies LI-COR IRDye 800CW Goat anti-Rabbit IgG (92532211, Licor, Lincoln, NE) and IRDye680RD goat anti-mouse IgG (925-68070, Li-Cor, Lincoln, NE). Image acquisition was performed using Li-Cor Odyssey System.

### Immunofluorescence

Doxycycline induced *Plcg1-*null fibroblasts expressing OptoHalo/OptoPLC-γ1 variants were plated on coverslips and allowed to spread overnight in high-glucose DMEM supplemented with 10% (v/v) FBS. Cells were labeled with 200 nM JF646 Halo ligand (GA1120, Promega, Madison, WI) per the manufacturer’s instructions, washed in warm phosphate buffer saline (PBS), fixed in 4% paraformaldehyde (PFA, Electron Microscopy Sciences, Hatfield, PA) in PBS for 15 min, and permeabilized with 0.1% Triton in PBS for 10 min. After blocking in 1% bovine serum albumin (BSA)/PBS for 1 hour, cells were incubated with anti-V5 (1:750) at 4C, overnight, followed by incubation with secondary antibodies RRX-conjugated-donkey anti-mouse (715-295-151, Jackson ImmunoResearch Laboratories, West Grove, PA) at room temperature for 1 hour. Coverslips were mounted to slides using the Prolong Glass antifade mountant (P36980, Thermo Fisher Scientific, Waltham, MA). Images were acquired using a Zeiss LSM800 confocal microscope equipped with an EC Plan-Neofluar 40x/1.30 oil DIC M27 objective lens.

### Live cell imaging and focal photoactivation

Live cell microscopy and focal photoactivation experiments were performed on a Zeiss LSM800, equipped with a humidity chamber, at 37°C and 5% CO_2_. Images were acquired with a 40x oil-immersion objective. For single ROI photoactivation experiments, cells were plated on glass-bottom dishes (Mattek Corporation, Ashland, MA) in high-glucose DMEM supplemented with 10% (v/v) FBS, induced with doxycycline (500 ng/mL for *Plcg1-*null fibroblasts or 1 µg/mL for JR20 fibrbolasts) overnight, and labeled with 200 nM JF646 Halo ligand per the manufacturer’s instructions. Cells with qualitatively comparable JF646 expression were photoactivated by illuminating an ROI, typically a 27 µm x 27 µm box encompassing the edge of the cell leaving space to maintain edge photoactivation should the cell protrude, for 8-10 seconds with a 10 mW 488 nm laser at 0.1% power (1x). Images were acquired every 15 seconds. For sustained migration experiments, an optotactic gradient was created by 20 µm x 20 µm adjacent ROIs, with stepwise increases of a 10 mW 488 nm laser power: ROI 1 encompassing the edge of the cell was set to 0.03% laser power (0.3x), ROI 2 was set to 0.06% laser power (0.6x), ROI 3 was set to 0.08 laser power (0.8x), ROI 4 was set to 0.1% laser power (1x), ROI 5 was set to 0.12% laser power (1.2x), ROI 6 was set to 0.14% laser power (1.4x), ROI 7 was set to 0.16% laser power (1.6x). Images were acquired every 30 seconds for an average of 16 hours. For calcium signaling experiments using the R-GECO biosensor, images were captured every 5 seconds for at least 2.5 minutes prior to photoactivation. Upon photoactivation, ROI illumination remained approximately continuous between frames. Photoactivation experiments including single pharmacological inhibition and a DMSO control were performed in the same dish. After photoactivating cells in the presence of DMSO, half of the liquid volume was replaced with inhibitor-containing media to achieve the target inhibitor concentration while maintaining the DMSO concentration. Cells were equilibrated with the inhibitor for at least 20 minutes prior to photoactivating individual cells. For photoactivation experiments during simultaneous calcium chelation and PKC inhibition, cells were plated in an 8-well, glass-bottom chamber to limit the duration of the double inhibition to one hour.

For double ROI experiments, Mattek dishes were coated with 1 µg/mL FN for 30 minutes at 37°C followed by a PEG-Poly-L-Lysine backfill (0.1 mg/mL) for 30 minutes at room temperature. Cells were then plated in high-glucose DMEM with 10% (v/v) FBS, induced with 1 µg/mL doxycycline overnight and labeled with 200 nM JFX554 per the manufacturer’s instructions. Cells with qualitatively comparable JFX554 expression were simultaneously photoactivated with two ROIs, both 60 µm wide by 40 µm tall rectangles, using a 10mW 488 nm laser. The ROIs were placed on the upper and lower edges of the cells, with as close to equal cell area within each ROI as possible. One of the ROIs was set to 0.01% laser power (0.1x), and the other to 0.02% (0.2x); the 0.2x ROI was usually chosen to be on the side of the cell that qualitatively appeared to be the trailing side pre-activation. Cells were photoactivated for approximately 25 seconds, with images acquired every 30 seconds. Simultaneous photoactivation lasted for 30 minutes and, when possible, a follow-up experiment was performed within 10 minutes of the end of the first in which the ROIs were repositioned, replacing the ROIs on the edges of the cell such that equal cell area was within each ROI and setting the 0.2x ROI on the retracting edge and the 0.1x ROI on the protruding edge of the cell.

To image lipid dynamics, doxycycline induced *Plcg1-*null fibroblasts expressing OptoHalo/OptoPLC-γ1 were transfected with C1A domain of PKCγ fused to mScarlet (DAG biosensor) and plated on glass-bottom dishes. Two 7 µm x 7 µm ROIs were positioned inside of the cell: ROI 1 was illuminated using a 10 mW 488 nm laser power at 0.1% (1x), while RO1 2 was not illuminated. Images were acquired every 10 seconds.

### Image quantification

Image analysis was performed using a combination of MATLAB (Mathworks) and Python. Fluorescent images were background subtracted and segmented via manual thresholding. The HaloTag fluorescence enrichment due to photoactivation was quantified by normalizing the ratio of the mean fluorescence intensity within the photoactivated ROI to that of the entire cell mask in each frame, relative to the average intensity across the ten frames preceding photoactivation. The resulting normalized intensity time series was fit to an exponential plateau function: 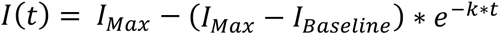, where 𝐼*_Max_* represents the approximate fold change in local intensity. To estimate the exponential decay rate of the normalized local intensity (i.e., dissociation rate) upon cessation of photoactivation, the ratio of the instantaneous slope to the normalized intensity was computed for the first three post-photoactivation frames and averaged.

To visualize the effect of photoactivation on cell morphology and motility, each frame’s cell mask was subtracted from the frame immediately preceding photoactivation, generating regions that denote protruded, retracted, and unchanged pixels. To spatially quantify area changes, we calculated the angle (rounded to the nearest degree) of each protruded/retracted pixel relative to the line connecting the ROI centroid to the cell centroid just before photoactivation. The ‘cell front’ and ‘cell rear’ were defined as 90° spans centered on 0° and 180°, respectively. The change in area was quantified by summing the pixel counts for all angles corresponding to these categorizations. Spatiotemporal maps of edge velocity were constructed as previously described(Welf et al., 2012; Johnson et al., 2015). Protruded/retracted pixels between consecutive frames were binned according to the aforementioned spatial reference and the sum of the pixel counts in each angular bin were converted to a velocity. Overlaid on the map is the angular position of the photoactivated ROI at the start of photoactivation. To quantify the effect of photoactivation on calcium signaling, the mean R-GECO fluorescence intensity of photoactivated cells was measured for each frame in the time series. The maximum R-GECO fluorescence intensity within the first 40 seconds of ROI illumination was then normalized to the average intensity of the preceding frames. To quantify the effect of photoactivation on lipid composition, the mean DAG biosensor fluorescence intensity was measured for each frame in an illuminated ROI and in a non-illuminated ROI in the time series. The DAG fluorescence intensity during photoactivation was normalized by the average intensity before photoactivation.

### Phospholipase activity assay

Two days prior to stimulation, cells harboring inducible OptoHalo or OptoPLC-γ1 variants and cultured in high-glucose DMEM containing 10% (v/v) FBS and 3 µg/mL puromycin were treated with 500 ng/mL doxycycline. The day before stimulation, cells were seeded in 6-well plates, with the same media composition, at ∼60,000 cells/well and allowed to spread. The spread cells were metabolically labeled overnight with [^3^H]myo-inositol (ART 0116A, American Radiolabeled Chemicals, Inc., St. Louis, MO) at 1 µCi/well in inositol-free DMEM containing the same additives as above. Cells stimulated with light were treated with assay buffer (20 mM HEPES (pH 7.4), 200 µg/mL fatty-acid free BSA, and 10 mM LiCl in Hank’s Balanced Salt Solution; all concentrations are final) and immediately subjected to global photoactivation, which was achieved by positioning plates 12 inches below an array of blue LED lights for 15 minutes. Matched dark samples were treated identically but without photoactivation. Cells challenged with PDGF were treated similarly to cells treated in the dark, except they were incubated with assay buffer containing 1 nM PDGF-BB (final concentration). All treatments were performed in a humified 37°C incubator with 5% CO2.

The accumulation of [^3^H]inositol phosphates was quantified as described previously, except that plates were protected from light during lysis (Waldo et al., 2010). Briefly, plates were transferred to ice, the medium was removed, and cells were lysed with ice-cold 50 mM formic acid for 45 minutes; during this time, plates were protected from light. Lysis was stopped by adding ice-cold 150 mM ammonium hydroxide. [^3^H]Inositol phosphates were captured from the soluble lysate using Dowex columns. The resin was washed with 50 mM ammonium formate, and bound [^3^H]inositol phosphates were eluted in 1.2 M ammonium formate containing 0.1 M formic acid. The amount of radioactivity in each sample was quantified by liquid scintillation counting.

## ACKNOWLEDGMENTS

We acknowledge Dr. Mitchell T. Butler, Dr. Bhagawat Subramanian, and Mark Hazelbaker for helpful discussions and tactical insight regarding cell line development, microscopy, and quantitative image analysis, as well as the ‘PLC Group’ for helpful discussions regarding PLC-γ1 structure and regulation. Research reported in this publication was supported by the National Institute of General Medical Sciences under award nos. R01 GM141691 (to J.M.H.) and R35 GM130312 (to J.E.B.). The content is solely the responsibility of the authors and does not necessarily represent the official views of the National Institutes of Health.

## AUTHOR CONTRIBUTIONS

RA: conceptualization, designed research, performed research, analyzed data, wrote and edited the paper; PFS: conceptualization, designed research, performed research, analyzed data, wrote and edited the paper; HT: designed research, performed research, analyzed data, wrote and edited the paper; NH: designed research, performed research, analyzed data, wrote and edited the paper; JS: conceptualization, designed research, supervision, funding acquisition, reviewed and edited the paper; JEB: conceptualization, designed research, supervision, funding acquisition, reviewed and edited the paper; JMH: conceptualization, designed research, supervision, funding acquisition, wrote and edited the paper.

## DECLARATION OF INTEREST

The authors declare no competing interests.

## SUPPLEMENTAL FIGURES

**Figure S1:**
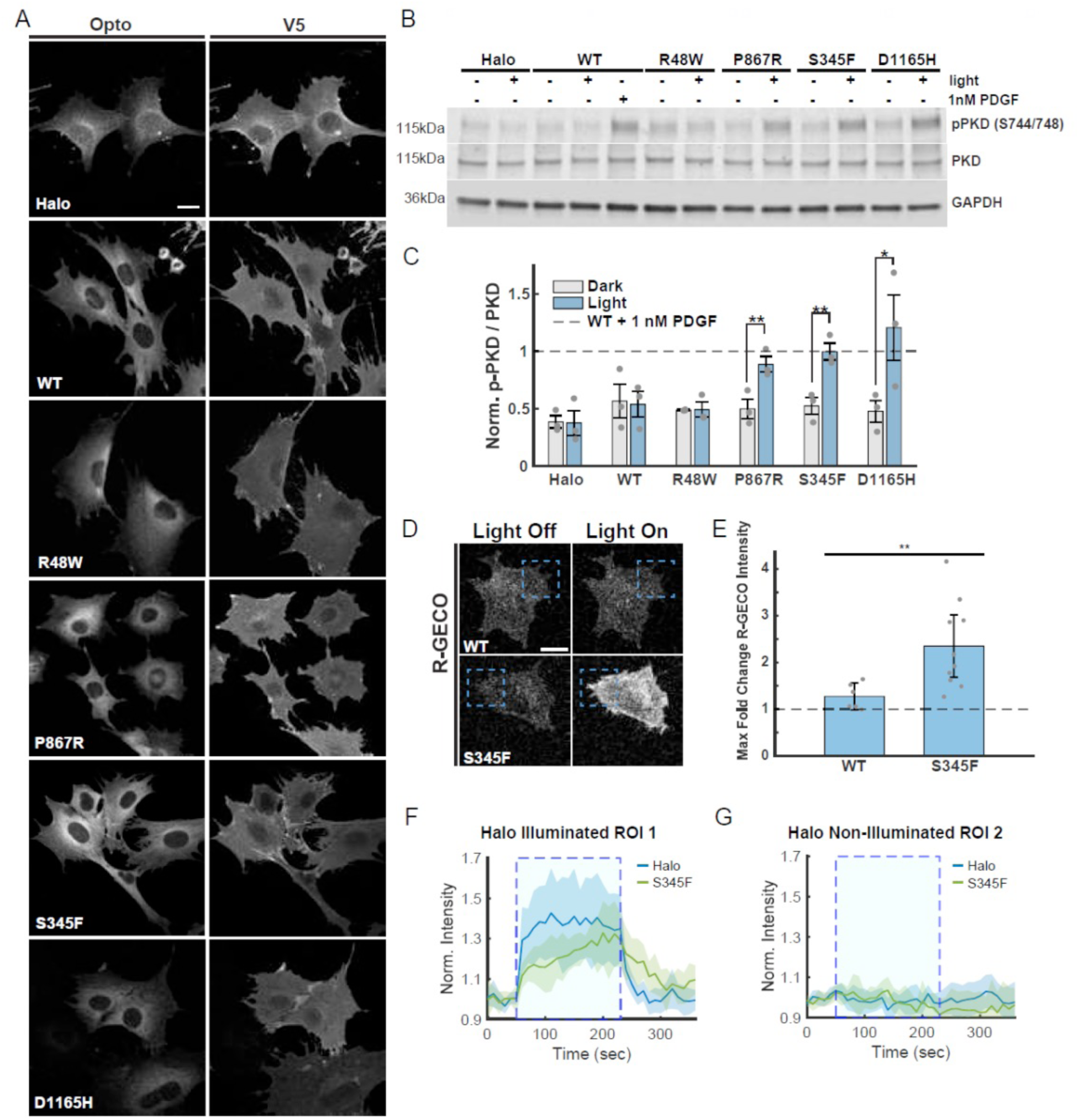
Inducible expression of OptoPLC-γ1 enables spatiotemporal control of membrane recruitment and enzyme activity. **(A)** Confocal, fluorescence micrographs of fixed, non-stimulated *Plcg1*-null fibroblasts induced to express V5-iLID-CaaX and Halo-SspBµ or PLCγ1-Halo-SspBµ variants. Cells were labeled with JF646 and immunostained for V5. **(B)** Immunoblot and **(C)** quantification of PKD and associated Ser744/748 phosphorylation in *Plcg1*-null fibroblasts rescued with either OptoHalo or OptoPLC-γ1 variants and globally stimulated with blue light, or 1 nM PDGF, for 5 minutes. Induction was performed by treating with 500 ng/ml doxycycline for 48 hours prior to stimulation. Blot is representative of n = 3 biological replicates. Phosphorylated S744/748 data are first normalized by total PKD and then by the p-PKD/PKD ratio of the WT OptoPLC-γ1 treated with 1 nM PDGF. The normalized data are reported as the mean. To test if normalized p-PKD increased upon light stimulation, dark vs. light treatments for each Opto variant were compared using a paired one-sided t-test. *p<0.05 and **p<0.01. **(D)** Representative live-cell confocal micrographs of the calcium biosensor R-GECO expressed in *Plcg1*-null fibroblasts rescued with OptoPLC-γ1 WT (n = 6 cells) or OptoPLC-γ1 S345F (n = 10 cells) and focally photoactivated. Dashed, blue box indicates region of interest (ROI) illuminated with a 488 nm laser. **(E)** Maximum fold change, relative to pre-photoactivation baseline, of whole-cell R-GECO fluorescence intensity observed up to 40 seconds after photoactivation. Data are reported as the mean ± 95% confidence interval. WT and S345F were compared using a two-sided Welch’s t-test. **p<0.01. **(F-G)** Time-course of Halo-JF646 enrichment within the illuminated ROI (ROI 1, blue) and non-illuminated ROI (ROI 2, black) in *Plcg1*-null fibroblasts expressing the aforementioned DAG biosensor and rescued with either OptoHalo (n = 8 cells) or OptoPLC-γ1 S345F (n = 7 cells). Solid, colored lines represent the mean of the respective intensities, and the shaded band regions indicate the 95% confidence interval.

**Figure S2:**
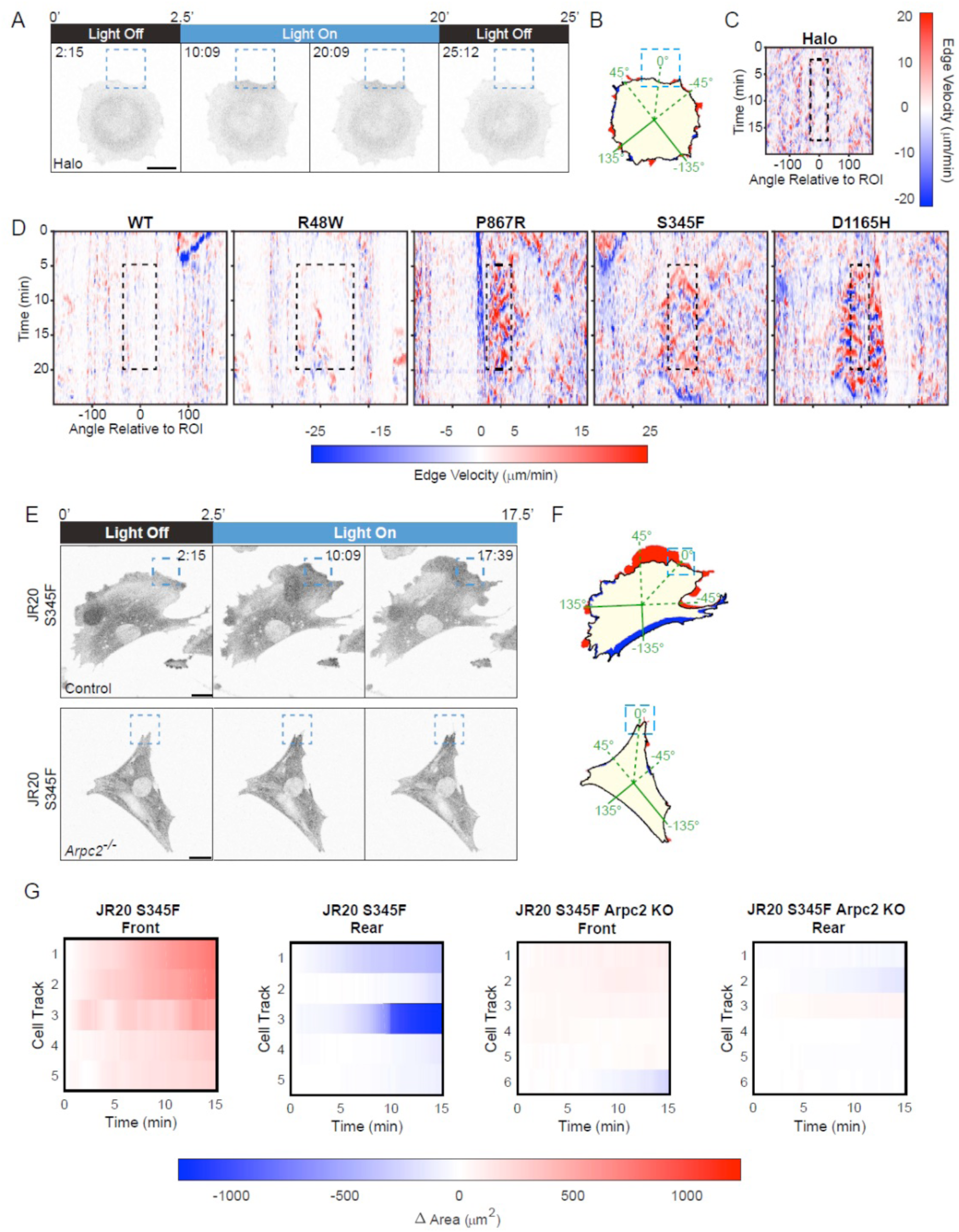
Photoactivating S345F produces protrusion within and proximal to the illuminated ROI and retraction on the opposing side of the cell. **(A)** Time-lapse confocal micrographs of *Plcg1*-null fibroblasts expressing OptoHalo labeled with JF646 ligand, and photoactivated as indicated. Dashed, blue box indicates ROI illuminated with a 488 nm laser. Time indicates min:sec and scale bars = 20 µm. **(B)** Net protrusion (red) and retraction (blue) for the representative cell shown in **A**. **(C)** Spatiotemporal map of edge velocity prior to, during, and after photoactivation for the representative OptoHalo cell shown in **A**. Zero degrees is defined by the line linking the centroid of the ROI and the centroid of the cell immediately prior to photoactivation (see pixel map in **B**). The dashed, black box denotes the spatiotemporal position of the illuminated ROI. **(D)** Spatiotemporal map of edge velocity prior to, during, and after photoactivation for the representative OptoPLC-γ1 variants shown in Fig. 2A**&B**. Here and hereafter, the dashed, black box denotes the spatiotemporal position of the ROI at the time of photoactivation. **(E)** Time-lapse confocal micrographs of control or tamoxifen-treated (*Arpc2^-/-^)* JR20 fibroblasts, expressing OptoPLC-γ1 S345F labeled with JF646 ligand, and photoactivated as indicated. Dashed, blue box indicates ROI illuminated with a 488 nm laser. Time indicates min:sec and scale bars = 20 µm. **(F)** Net protrusion (red) and retraction (blue) for the representative cell shown in **E**. **(G)** Heatmaps noting the change in area at the ‘cell front’ and ‘cell rear’ with respect to time for each photoactivated control or tamoxifen-treated JR20 fibroblasts expressing OptoPLC-γ1 S345F. The ‘cell front’ is defined as the 90-degree window centered on the line between the ROI centroid and cell centroid immediately prior to photoactivation as depicted by the cartoon (right). The ‘cell rear’ is defined as the 90-degree window mirroring the ‘cell front’. Time is scaled relative to the start of photoactivation.

**Figure S3:**
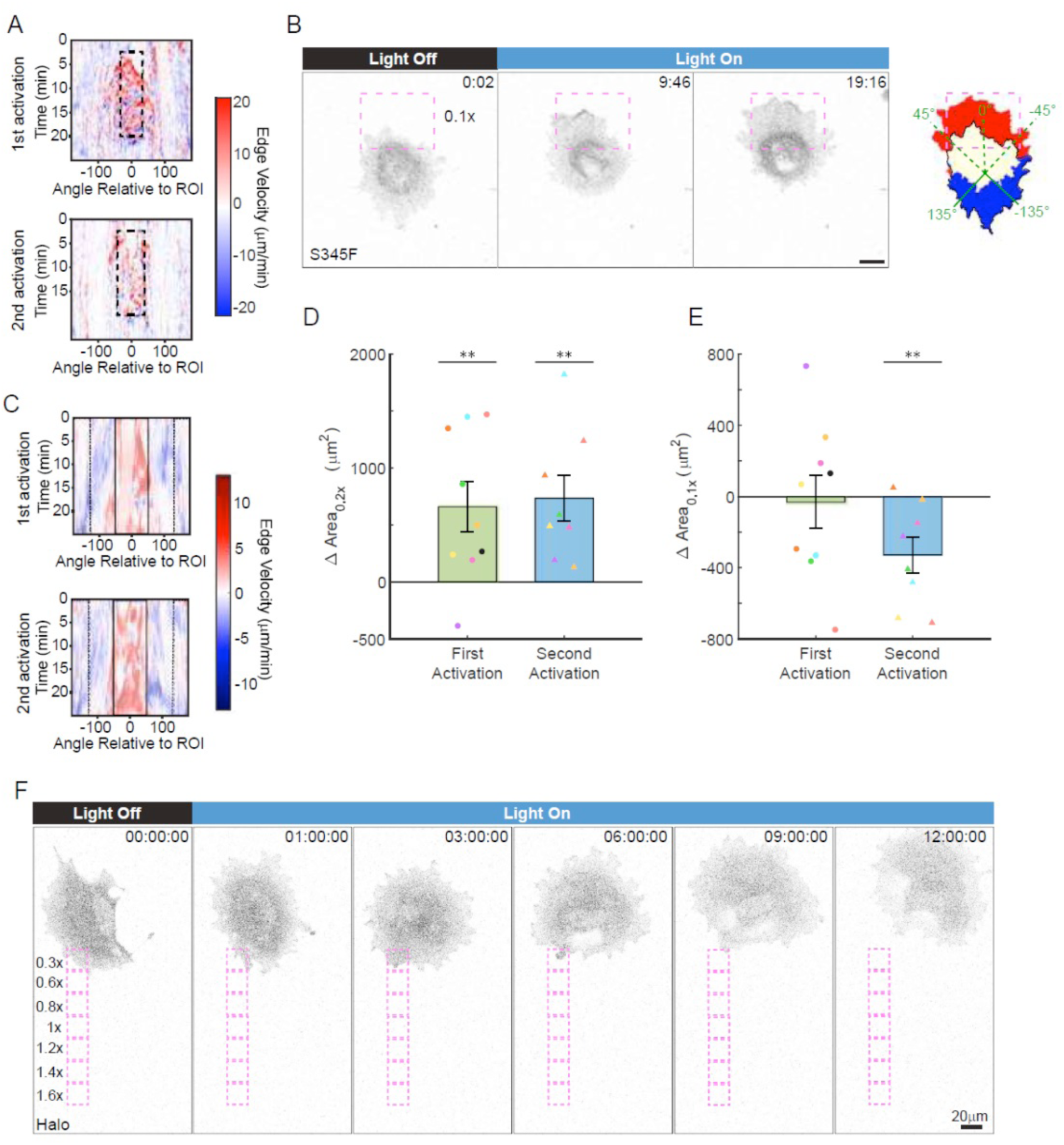
S345F photoactivation at 0.1x relative power polarizes cell motility. **(A)** Spatiotemporal map of edge velocity prior to, during, and after photoactivation for the first (top) and second (bottom) photoactivations of the cell shown in Fig. 4A**&B**. The dashed, black box denotes the spatiotemporal position of the illuminated ROI. **(B)** Time-lapse confocal micrographs, along with net protrusion (red) and retraction (blue), of an OptoPLC-γ1 S345F cell labeled with JFX554 ligand, photoactivated as indicated. Dashed, pink box indicates ROI illuminated with a 488-nm laser set to 0.01% (0.1x) power. Time indicates min:sec, and scale bar = 20 µm. **(C)** Spatiotemporal map of edge velocity prior to, during, and after photoactivation for the first (top) and second (bottom) photoactivations of the cell shown in Fig. 4C**&D**. Black boxes denote the spatiotemporal positions of 0.2x (solid) and 0.1x (dashed) ROIs. **(D-E)** Quantification of change in area within and proximal to the 0.2x (**D**) and 0.1x (**E**) ROIs for cells photoactivated under the dual ROI and reversion protocol. Multiple photoactivations of each cell are represented in all plots with the same color. Quantifications in (**D&E**) include 8 cells photoactivated twice sequentially (30-minute duration each, <10 minutes between) and a single cell, colored in black, photoactivated only once. Data are presented as the mean ± SEM. Delta area quantifications were compared against zero by the appropriate (right-tailed for **D**, left-tailed for **E**) one-tailed t-tests. **: p < 0.01 and *: p < 0.05. **(F)** Time-lapse confocal micrograph of a cell expressing OptoHalo labeled with JF646 ligand and photoactivated as indicated. Time indicates h:min:sec, and scale bar = 20 µm. Representative of n = 2 cells.

**Figure S4:**
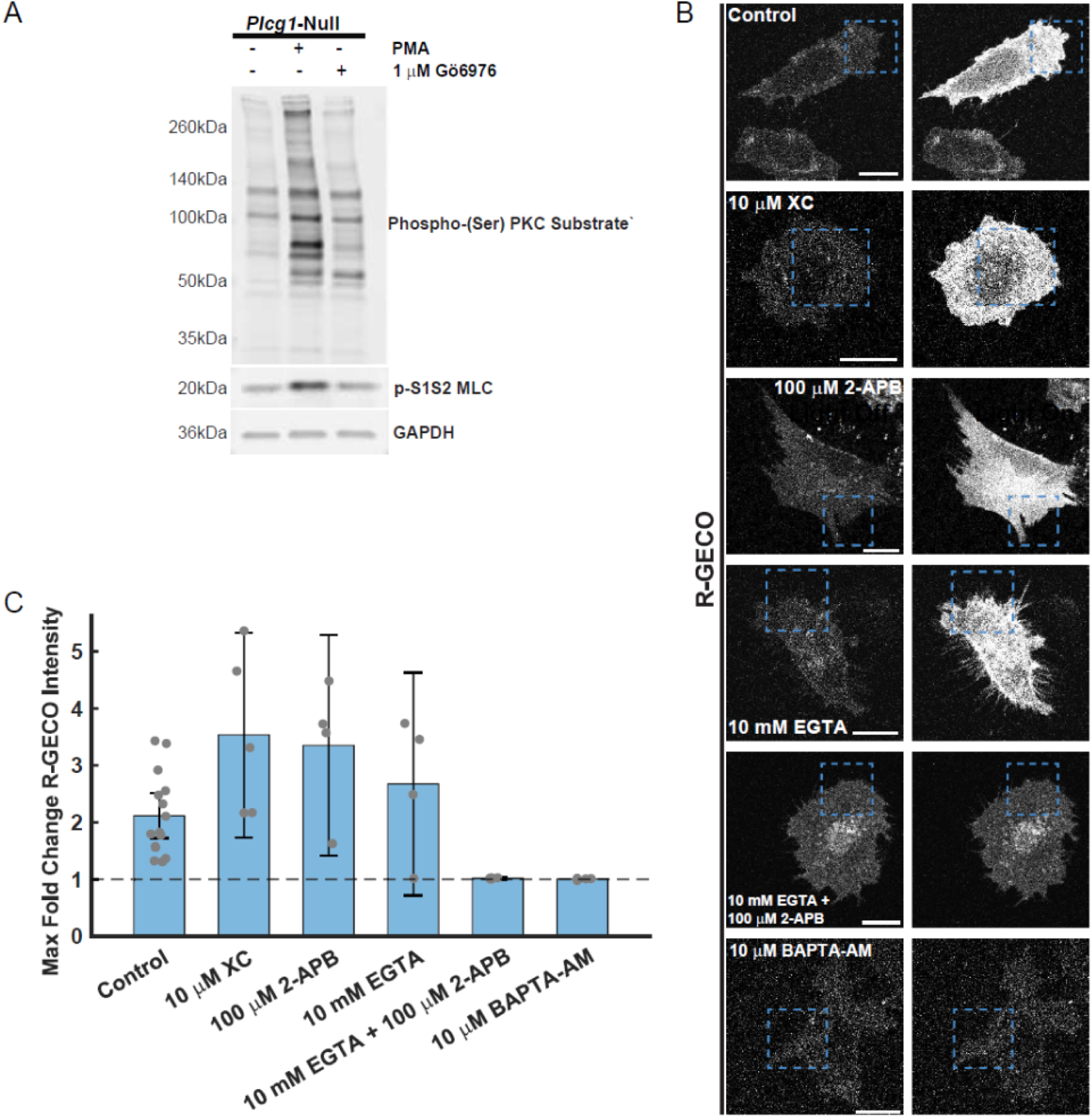
Validation of pharmacological inhibition of PKCα and calcium signaling. **(A)** Immunoblot of serum-starved *Plcg1*-null fibroblasts stimulated for 5 minutes with 200 nM PMA in the presence of either a vehicle control (0.1% DMSO) or Gӧ6976 and subsequently blotted with antibodies against phospho-(ser) PKC substrate and phosphorylated Ser1Ser2 myosin light chain (MLC). Blot shows the result one biological replicate. **(B)** Representative live-cell confocal micrographs of the calcium biosensor R-GECO expressed in *Plcg1*-null fibroblasts rescued with OptoPLC-γ1 S345F and focally photoactivated without or in the presence of the listed treatments (control: n =15 cells, 10 µM XC: n = 5 cells, 100 µM 2-APB: n = 4 cells, 10 mM EGTA: n = 4 cells, 10 mM EGTA + 100 µM 2-APB: n = 4 cells, 10 µM BAPTA-AM: n = 4 cells). Dashed, blue box indicates region of interest (ROI) illuminated with a 488 nm laser. **(C)** Maximum fold change, relative to pre-photoactivation baseline, of whole-cell R-GECO fluorescence intensity observed up to 40 seconds after photoactivation. Data are reported as the mean ± 95% confidence interval.

**Figure S5:**
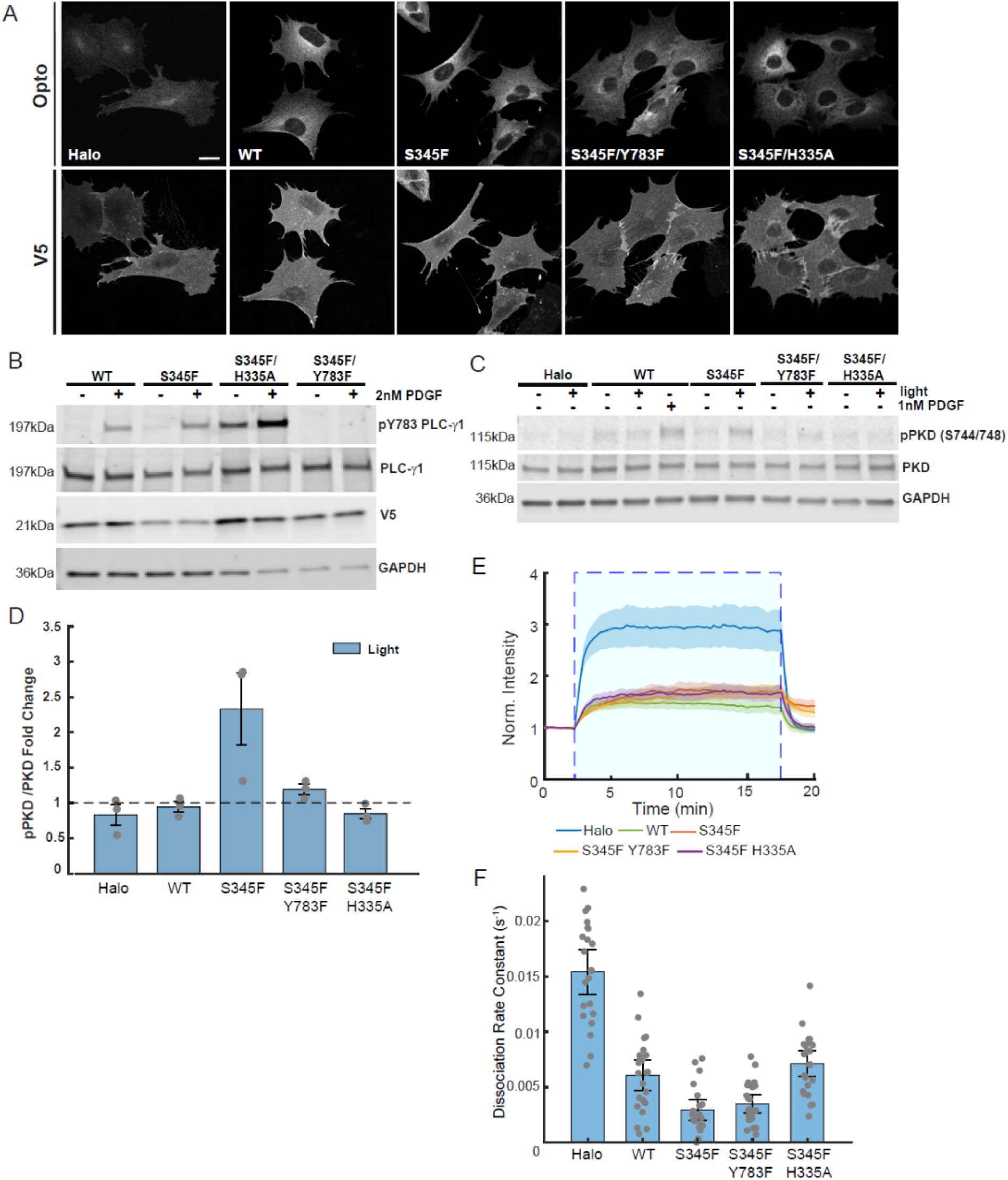
OptoPLC-γ1 S345F double-mutants are inducibly expressed with comparable V5/Halo ratios as WT and S345F variants. **(A)** Confocal, fluorescence micrographs of fixed, non-stimulated *Plcg1*-null fibroblasts induced to express V5-iLID-CaaX and Halo-SspBµ or PLCγ1-Halo-SspBµ variants. Cells were labeled with JF646 and immunostained for V5. **(B)** Immunoblot of PLC-γ1 and associated Tyr783 phosphorylation in *Plcg1*-null fibroblasts rescued with the listed OptoPLC variants and stimulated with 2 nM PDGF, for 5 minutes. Induction was performed by treating with 500 ng/ml doxycycline for 48 hours prior to stimulation. Blot is representative of a single biological replicate. **(C)** Immunoblot of PKD and associated Ser744/748 phosphorylation in *Plcg1*-null fibroblasts rescued with either OptoHalo or OptoPLC variants and globally stimulated with blue light, or 1 nM PDGF, for 5 minutes. Induction was performed by treating with 500 ng/ml doxycycline for 48 hours prior to stimulation. Blot is representative of n = 3 biological replicates. **(D)** Immunoblot quantification of PKD S744/748 phosphorylation across OptoHalo/OptoPLC variants globally stimulated with light for 5 minutes. Total and phospho-ser 744/748 PKD are first normalized by their associated GAPDH loading controls before calculating the ratio of phospho-PKD to total PKD. The light condition is then normalized by the dark state for each variant. Data reported as the mean ±95% confidence interval. **(E)** Time-course of Halo-JF646 enrichment within the photoactivated ROI in *Plcg1*-null fibroblasts rescued with the indicated OptoHalo/OptoPLC-γ1 variants upon focal illumination with a 488 nm laser. Solid, colored lines represent the mean of the respective intensities for OptoHalo (n = 22 cells), OptoPLC-γ1 WT (n = 23 cells), OptoPLC-γ1 S345F (n = 22 cells), OptoPLC-γ1 S345F/Y783F (n = 23 cells), and OptoPLC-γ1 S345F/H335A (n = 24 cells). The shaded band regions indicate the 95% confidence interval. **(F)** Estimated decay rate constants upon stopping focal activation (see Materials and Methods). Data are presented as the mean ± 95% confidence interval.

## Notes

### Competing Interest Statement

The authors have declared no competing interest.

### Summary of Updates

In response to review comments, the major changes are: Using a translocation biosensor that binds diacylglycerol (DAG) at the plasma membrane, we confirmed that photo-stimulation of OptoPLC-γ1 S345F activates PIP2 hydrolysis locally, with kinetics that mirror the membrane recruitment of the enzyme (current Fig. 2D&E and Suppl. Fig. S1F&G). Using the previously described JR20 conditional Arpc2 knockout line, we show that the light-induced motility response in fibroblasts expressing OptoPLC-γ1 S345F is dependent on the Arp2/3 complex (current Suppl. Fig. S2E-G); apparently, protrusion is achieved by mobilizing the lamellipodial F-actin machinery. Most importantly, we now demonstrate that a staircase gradient of light intensity directs persistent migration of cells expressing OptoPLC-γ1 S345F (current Fig. 4F/Video 4) but not of control cells (current Suppl. Fig. S3F/Video 5). This result is reflected in our new title, Optogenetic control of PLC-γ1 activity directs cell motility, and in the abstract. We tested the effect of combining the PKC and calcium inhibitor perturbations (1 microM Go6976 and 10 microM BAPTA-AM) and found that the effect - partial inhibition but not ablation of the polarization response - is qualitatively the same as that of either inhibitor alone (current Fig. 7).

## REFERENCES

Albert, A.P., S.N. Saleh, and W.A. Large. 2008. Inhibition of native TRPC6 channel activity by phosphatidylinositol 4,5-bisphosphate in mesenteric artery myocytes. J. Physiol. 586:3087–3095. doi:10.1113/jphysiol.2008.153676.

Asokan, S.B., H.E. Johnson, A. Rahman, S.J. King, J.D. Rotty, I.P. Lebedeva, J.M. Haugh, and J.E. Bear. 2014. Mesenchymal chemotaxis requires selective inactivation of myosin II at the leading edge via a noncanonical PLCγ/PKCα pathway. Dev. Cell. 31:747–760. doi:10.1016/j.devcel.2014.10.024.

Barger, C.J., C. Branick, L. Chee, and A.R. Karpf. 2019. Pan-cancer analyses reveal genomic features of FOXM1 overexpression in cancer. Cancers. 11:251. doi:10.3390/cancers11020251.

Bear, J.E., and J.M. Haugh. 2014. Directed migration of mesenchymal cells: where signaling and the cytoskeleton meet. Curr. Opin. Cell Biol. 30:74–82. doi:10.1016/j.ceb.2014.06.005.

Bisaria, A., A. Hayer, D. Garbett, D. Cohen, and T. Meyer. 2020. Membrane-proximal F-actin restricts local membrane protrusions and directs cell migration. Science. 368:1205–1210. doi:10.1126/science.aay7794.

Black, A.R., and J.D. Black. 2021. The complexities of PKCα signaling in cancer. Adv. Biol. Regul. 80:100769. doi:10.1016/j.jbior.2020.100769.

Bouma, G., S.O. Burns, and A.J. Thrasher. 2009. Wiskott-Aldrich Syndrome: Immunodeficiency resulting from defective cell migration and impaired immunostimulatory activation. Immunobiology. 214:778–790. doi:10.1016/j.imbio.2009.06.009.

Braiman, A., M. Barda-Saad, C.L. Sommers, and L.E. Samelson. 2006. Recruitment and activation of PLCgamma1 in T cells: a new insight into old domains. EMBO J. 25:774–784. doi:10.1038/sj.emboj.7600978.

Bravo-Cordero, J.J., L. Hodgson, and J. Condeelis. 2012. Directed cell invasion and migration during metastasis. Curr. Opin. Cell Biol. 24:277–283. doi:10.1016/j.ceb.2011.12.004.

Cai, L., T.W. Marshall, A.C. Uetrecht, D.A. Schafer, and J.E. Bear. 2007. Coronin 1B coordinates Arp2/3 complex and cofilin activities at the leading edge. Cell. 128:915–929. doi:10.1016/j.cell.2007.01.031.

Chandra, A., M.T. Butler, J.E. Bear, and J.M. Haugh. 2022. Modeling cell protrusion predicts how myosin II and actin turnover affect adhesion-based signaling. Biophys. J. 121:102–118. doi:10.1016/j.bpj.2021.11.2889.

Chen, D., and M. Simons. 2021. Emerging roles of PLCγ1 in endothelial biology. Sci. Signal. 14:eabc6612. doi:10.1126/scisignal.abc6612.

Clapham, D.E. 2007. Calcium Signaling. Cell. 131:1047–1058. doi:10.1016/j.cell.2007.11.028.

Cooke, M., and M.G. Kazanietz. 2022. Overarching roles of diacylglycerol signaling in cancer development and antitumor immunity. Sci. Signal. 15:eabo0264. doi:10.1126/scisignal.abo0264.

De Belly, H., and O.D. Weiner. 2024. Follow the flow: Actin and membrane act as an integrated system to globally coordinate cell shape and movement. Curr. Opin. Cell Biol. 89:102392. doi:10.1016/j.ceb.2024.102392.

DeBell, K.E., B.A. Stoica, M.C. Verí, A. Di Baldassarre, S. Miscia, L.J. Graham, B.L. Rellahan, M. Ishiai, T. Kurosaki, and E. Bonvini. 1999. Functional independence and interdependence of the Src homology domains of phospholipase C-gamma1 in B-cell receptor signal transduction. Mol. Cell. Biol. 19:7388–7398. doi:10.1128/MCB.19.11.7388.

Encell, L.P., R. Friedman Ohana, K. Zimmerman, P. Otto, G. Vidugiris, M.G. Wood, G.V. Los, M.G. McDougall, C. Zimprich, N. Karassina, R.D. Learish, R. Hurst, J. Hartnett, S. Wheeler, P. Stecha, J. English, K. Zhao, J. Mendez, H.A. Benink, N. Murphy, D.L. Daniels, M.R. Slater, M. Urh, A. Darzins, D.H. Klaubert, R.F. Bulleit, and K.V. Wood. 2012. Development of a dehalogenase-based protein fusion tag capable of rapid, selective and covalent attachment to customizable ligands. Curr. Chem. Genomics. 6:55–71. doi:10.2174/1875397301206010055.

Ferguson, G.J., L. Milne, S. Kulkarni, T. Sasaki, S. Walker, S. Andrews, T. Crabbe, P. Finan, G. Jones, S. Jackson, M. Camps, C. Rommel, M. Wymann, E. Hirsch, P. Hawkins, and L. Stephens. 2007. PI(3)Kγ has an important context-dependent role in neutrophil chemokinesis. Nat. Cell Biol. 9:86–91. doi:10.1038/ncb1517.

Frei, M.S., M. Tarnawski, M.J. Roberti, B. Koch, J. Hiblot, and K. Johnsson. 2022. Engineered HaloTag variants for fluorescence lifetime multiplexing. Nat. Methods. 19:65–70. doi:10.1038/s41592-021-01341-x.

Gafni, J., J.A. Munsch, T.H. Lam, M.C. Catlin, L.G. Costa, T.F. Molinski, and I.N. Pessah. 1997. Xestospongins: potent membrane permeable blockers of the inositol 1,4,5-trisphosphate receptor. Neuron. 19:723–733. doi:10.1016/s0896-6273(00)80384-0.

Gambardella, J., M.B. Morelli, X. Wang, V. Castellanos, P. Mone, and G. Santulli. 2021. The discovery and development of IP3 receptor modulators: an update. Expert Opin. Drug Discov. 16:709–718. doi:10.1080/17460441.2021.1858792.

Gresset, A., S.N. Hicks, T.K. Harden, and J. Sondek. 2010. Mechanism of phosphorylation- induced activation of phospholipase C-gamma isozymes. J. Biol. Chem. 285:35836– 35847. doi:10.1074/jbc.M110.166512.

Gresset, A., J. Sondek, and T.K. Harden. 2012. The phospholipase C isozymes and their regulation. Subcell. Biochem. 58:61–94. doi:10.1007/978-94-007-3012-0_3.

Guarnieri, F.C., A. de Chevigny, A. Falace, and C. Cardoso. 2018. Disorders of neurogenesis and cortical development. Dialogues Clin. Neurosci. 20:255–266. doi:10.31887/DCNS.2018.20.4/ccardoso.

Guntas, G., R.A. Hallett, S.P. Zimmerman, T. Williams, H. Yumerefendi, J.E. Bear, and B. Kuhlman. 2015. Engineering an improved light-induced dimer (iLID) for controlling the localization and activity of signaling proteins. Proc. Natl. Acad. Sci. U. S. A. 112:112–117. doi:10.1073/pnas.1417910112.

Hajicek, N., T.H. Charpentier, J.R. Rush, T.K. Harden, and J. Sondek. 2013. Autoinhibition and phosphorylation-induced activation of phospholipase C-γ isozymes. Biochemistry. 52:4810–4819. doi:10.1021/bi400433b.

Hajicek, N., N.C. Keith, E. Siraliev-Perez, B.R. Temple, W. Huang, Q. Zhang, T.K. Harden, and J. Sondek. 2019. Structural basis for the activation of PLC-γ isozymes by phosphorylation and cancer-associated mutations. eLife. 8:e51700. doi:10.7554/eLife.51700.

Hallett, R.A., S.P. Zimmerman, H. Yumerefendi, J.E. Bear, and B. Kuhlman. 2016. Correlating *in vitro* and *in vivo* activities of light-inducible dimers: a cellular optogenetics guide. ACS Synth. Biol. 5:53–64. doi:10.1021/acssynbio.5b00119.

Hao, J.-J., Y. Liu, M. Kruhlak, K.E. Debell, B.L. Rellahan, and S. Shaw. 2009. Phospholipase C-mediated hydrolysis of PIP2 releases ERM proteins from lymphocyte membrane. J. Cell Biol. 184:451–462. doi:10.1083/jcb.200807047.

Haugh, J.M., F. Codazzi, M. Teruel, and T. Meyer. 2000. Spatial sensing in fibroblasts mediated by 3ʹ phosphoinositides. J. Cell Biol. 151:1269–1280. doi:10.1083/jcb.151.6.1269.

Hennecke, M., M. Kwissa, K. Metzger, A. Oumard, A. Kröger, R. Schirmbeck, J. Reimann, and H. Hauser. 2001. Composition and arrangement of genes define the strength of IRES-driven translation in bicistronic mRNAs. Nucleic Acids Res. 29:3327–3334. doi:10.1093/nar/29.16.3327.

Hofmann, T., A.G. Obukhov, M. Schaefer, C. Harteneck, T. Gudermann, and G. Schultz. 1999. Direct activation of human TRPC6 and TRPC3 channels by diacylglycerol. Nature. 397:259–263. doi:10.1038/16711.

Hou, X., C. Ren, J. Jin, Y. Chen, X. Lyu, K. Bi, N.D. Carrillo, V.L. Cryns, R.A. Anderson, J. Sun, and M. Chen. 2025. Phosphoinositide signalling in cell motility and adhesion. Nat. Cell Biol. 27:736–748. doi:10.1038/s41556-025-01647-4.

Idevall-Hagren, O., E.J. Dickson, B. Hille, D.K. Toomre, and P. De Camilli. 2012. Optogenetic control of phosphoinositide metabolism. Proc. Natl. Acad. Sci. U. S. A. 109:E2316–2323. doi:10.1073/pnas.1211305109.

Jang, H.-J., P.-G. Suh, Y.J. Lee, K.J. Shin, L. Cocco, and Y.C. Chae. 2018. PLCγ1: Potential arbitrator of cancer progression. Adv. Biol. Regul. 67:179–189. doi:10.1016/j.jbior.2017.11.003.

Ji, Q.S., G.E. Winnier, K.D. Niswender, D. Horstman, R. Wisdom, M.A. Magnuson, and G. Carpenter. 1997. Essential role of the tyrosine kinase substrate phospholipase C-gamma1 in mammalian growth and development. Proc. Natl. Acad. Sci. U. S. A. 94:2999–3003. doi:10.1073/pnas.94.7.2999.

Johnson, H.E., S.J. King, S.B. Asokan, J.D. Rotty, J.E. Bear, and J.M. Haugh. 2015. F-actin bundles direct the initiation and orientation of lamellipodia through adhesion-based signaling. J. Cell Biol. 208:443–455. doi:10.1083/jcb.201406102.

Jones, N.P., and M. Katan. 2007. Role of phospholipase Cgamma1 in cell spreading requires association with a beta-Pix/GIT1-containing complex, leading to activation of Cdc42 and Rac1. Mol. Cell. Biol. 27:5790–5805. doi:10.1128/MCB.00778-07.

Jones, N.P., J. Peak, S. Brader, S.A. Eccles, and M. Katan. 2005. PLCγ1 is essential for early events in integrin signalling required for cell motility. J. Cell Sci. 118:2695–2706. doi:10.1242/jcs.02374.

Kataoka, K., Y. Nagata, A. Kitanaka, Y. Shiraishi, T. Shimamura, J. Yasunaga, Y. Totoki, K. Chiba, A. Sato-Otsubo, G. Nagae, R. Ishii, S. Muto, S. Kotani, Y. Watatani, J. Takeda, M. Sanada, H. Tanaka, H. Suzuki, Y. Sato, Y. Shiozawa, T. Yoshizato, K. Yoshida, H. Makishima, M. Iwanaga, G. Ma, K. Nosaka, M. Hishizawa, H. Itonaga, Y. Imaizumi, W. Munakata, H. Ogasawara, T. Sato, K. Sasai, K. Muramoto, M. Penova, T. Kawaguchi, H. Nakamura, N. Hama, K. Shide, Y. Kubuki, T. Hidaka, T. Kameda, T. Nakamaki, K. Ishiyama, S. Miyawaki, S.-S. Yoon, K. Tobinai, Y. Miyazaki, A. Takaori-Kondo, F. Matsuda, K. Takeuchi, O. Nureki, H. Aburatani, T. Watanabe, T. Shibata, M. Matsuoka, S. Miyano, K. Shimoda, and S. Ogawa. 2015. Integrated molecular analysis of adult T cell leukemia/lymphoma. Nat. Genet. 47:1304–1315. doi:10.1038/ng.3415.

Kim, H.K., J.W. Kim, A. Zilberstein, B. Margolis, J.G. Kim, J. Schlessinger, and S.G. Rhee. 1991. PDGF stimulation of inositol phospholipid hydrolysis requires PLC-gamma 1 phosphorylation on tyrosine residues 783 and 1254. Cell. 65:435–441. doi:10.1016/0092-8674(91)90461-7.

Kim, J.M., M. Lee, N. Kim, and W.D. Heo. 2016. Optogenetic toolkit reveals the role of Ca^2+^ sparklets in coordinated cell migration. Proc. Natl. Acad. Sci. 113:5952–5957. doi:10.1073/pnas.1518412113.

Kim, Y.-J., S. Tohyama, T. Nagashima, M. Nagase, Y. Hida, S. Hamada, A.M. Watabe, and T. Ohtsuka. 2024. A light-controlled phospholipase C for imaging of lipid dynamics and controlling neural plasticity. Cell Chem. Biol. 31:1336–1348.e7. doi:10.1016/j.chembiol.2024.03.001.

Krause, M., and A. Gautreau. 2014. Steering cell migration: lamellipodium dynamics and the regulation of directional persistence. Nat. Rev. Mol. Cell Biol. 15:577–590. doi:10.1038/nrm3861.

Lattanzio, R., M. Piantelli, and M. Falasca. 2013. Role of phospholipase C in cell invasion and metastasis. Adv. Biol. Regul. 53:309–318. doi:10.1016/j.jbior.2013.07.006.

Le Huray, K.I.P., T.D. Bunney, N. Pinotsis, A.C. Kalli, and M. Katan. 2022. Characterization of the membrane interactions of phospholipase Cγ reveals key features of the active enzyme. Sci. Adv. 8:eabp9688. doi:10.1126/sciadv.abp9688.

Leithner, A., A. Eichner, J. Müller, A. Reversat, M. Brown, J. Schwarz, J. Merrin, D.J.J. De Gorter, F. Schur, J. Bayerl, I. De Vries, S. Wieser, R. Hauschild, F.P.L. Lai, M. Moser, D. Kerjaschki, K. Rottner, J.V. Small, T.E.B. Stradal, and M. Sixt. 2016. Diversified actin protrusions promote environmental exploration but are dispensable for locomotion of leukocytes. Nat. Cell Biol. 18:1253–1259. doi:10.1038/ncb3426.

Mandal, S., S. Bandyopadhyay, K. Tyagi, and A. Roy. 2021. Recent advances in understanding the molecular role of phosphoinositide-specific phospholipase C gamma 1 as an emerging onco-driver and novel therapeutic target in human carcinogenesis. Biochim. Biophys. Acta Rev. Cancer. 1876:188619. doi:10.1016/j.bbcan.2021.188619.

Martiny-Baron, G., M.G. Kazanietz, H. Mischak, P.M. Blumberg, G. Kochs, H. Hug, D. Marmé, and C. Schächtele. 1993. Selective inhibition of protein kinase C isozymes by the indolocarbazole Gö 6976. J. Biol. Chem. 268:9194–9197.

Maruyama, T., T. Kanaji, S. Nakade, T. Kanno, and K. Mikoshiba. 1997. 2APB, 2- aminoethoxydiphenyl borate, a membrane-penetrable modulator of Ins(1,4,5)P3- induced Ca2+ release. J. Biochem. (Tokyo). 122:498–505. doi:10.1093/oxfordjournals.jbchem.a021780.

Melvin, A.T., E.S. Welf, Y. Wang, D.J. Irvine, and J.M. Haugh. 2011. In chemotaxing fibroblasts, both high-fidelity and weakly biased cell movements track the localization of PI3K signaling. Biophys. J. 100:1893–1901. doi:10.1016/j.bpj.2011.02.047.

Monypenny, J., D. Zicha, C. Higashida, F. Oceguera-Yanez, S. Narumiya, and N. Watanabe. 2009. Cdc42 and Rac Family GTPases regulate mode and speed but not direction of primary fibroblast migration during platelet-derived growth factor-dependent chemotaxis. Mol. Cell. Biol. 29:2730–2747. doi:10.1128/MCB.01285-08.

Nosbisch, J.L., J.E. Bear, and J.M. Haugh. 2022. A kinetic model of phospholipase C-γ1 linking structure-based insights to dynamics of enzyme autoinhibition and activation. J. Biol. Chem. 298:101886. doi:10.1016/j.jbc.2022.101886.

Raucher, D., T. Stauffer, W. Chen, K. Shen, S. Guo, J.D. York, M.P. Sheetz, and T. Meyer. 2000. Phosphatidylinositol 4,5-bisphosphate functions as a second messenger that regulates cytoskeleton-plasma membrane adhesion. Cell. 100:221–228. doi:10.1016/s0092-8674(00)81560-3.

van Rheenen, J., X. Song, W. van Roosmalen, M. Cammer, X. Chen, V. Desmarais, S.-C. Yip, J.M. Backer, R.J. Eddy, and J.S. Condeelis. 2007. EGF-induced PIP2 hydrolysis releases and activates cofilin locally in carcinoma cells. J. Cell Biol. 179:1247–1259. doi:10.1083/jcb.200706206.

Ridley, A.J., M.A. Schwartz, K. Burridge, R.A. Firtel, M.H. Ginsberg, G. Borisy, J.T. Parsons, and A.R. Horwitz. 2003. Cell migration: integrating signals from front to back. Science. 302:1704–1709. doi:10.1126/science.1092053.

Rotin, D., B. Margolis, M. Mohammadi, R.J. Daly, G. Daum, N. Li, E.H. Fischer, W.H. Burgess, A. Ullrich, and J. Schlessinger. 1992. SH2 domains prevent tyrosine dephosphorylation of the EGF receptor: identification of Tyr992 as the high-affinity binding site for SH2 domains of phospholipase C gamma. EMBO J. 11:559–567. doi:10.1002/j.1460-2075.1992.tb05087.x.

Rotty, J.D., C. Wu, E.M. Haynes, C. Suarez, J.D. Winkelman, H.E. Johnson, J.M. Haugh, D.R. Kovar, and J.E. Bear. 2015. Profilin-1 Serves as a Gatekeeper for Actin Assembly by Arp2/3- Dependent and -Independent Pathways. Dev. Cell. 32:54–67. doi:10.1016/j.devcel.2014.10.026.

Sadhu, R.K., A. Iglič, and N.S. Gov. 2023. A minimal cell model for lamellipodia-based cellular dynamics and migration. J. Cell Sci. 136:jcs260744. doi:10.1242/jcs.260744.

Sala, G., F. Dituri, C. Raimondi, S. Previdi, T. Maffucci, M. Mazzoletti, C. Rossi, M. Iezzi, R. Lattanzio, M. Piantelli, S. Iacobelli, M. Broggini, and M. Falasca. 2008. Phospholipase Cgamma1 is required for metastasis development and progression. Cancer Res. 68:10187–10196. doi:10.1158/0008-5472.CAN-08-1181.

Schuhmacher, M., A.T. Grasskamp, P. Barahtjan, N. Wagner, B. Lombardot, J.S. Schuhmacher, P. Sala, A. Lohmann, I. Henry, A. Shevchenko, Ü. Coskun, A.M. Walter, and A. Nadler. 2020. Live-cell lipid biochemistry reveals a role of diacylglycerol side-chain composition for cellular lipid dynamics and protein affinities. Proc. Natl. Acad. Sci. U. S. A. 117:7729– 7738. doi:10.1073/pnas.1912684117.

SenGupta, S., C.A. Parent, and J.E. Bear. 2021. The principles of directed cell migration. Nat. Rev. Mol. Cell Biol. 22:529–547. doi:10.1038/s41580-021-00366-6.

Senju, Y., M. Kalimeri, E.V. Koskela, P. Somerharju, H. Zhao, I. Vattulainen, and P. Lappalainen. 2017. Mechanistic principles underlying regulation of the actin cytoskeleton by phosphoinositides. Proc. Natl. Acad. Sci. U. S. A. 114:E8977–E8986. doi:10.1073/pnas.1705032114.

Servant, G., O.D. Weiner, P. Herzmark, T. Balla, J.W. Sedat, and H.R. Bourne. 2000. Polarization of chemoattractant receptor signaling during neutrophil chemotaxis. Science. 287:1037– 1040. doi:10.1126/science.287.5455.1037.

Siraliev-Perez, E., J.T. Stariha, R.M. Hoffmann, B.R. Temple, Q. Zhang, N. Hajicek, M.L. Jenkins, J.E. Burke, and J. Sondek. 2022. Dynamics of allosteric regulation of the phospholipase C- γ isozymes upon recruitment to membranes. eLife. 11:e77809. doi:10.7554/eLife.77809.

Tao, P., X. Han, Q. Wang, S. Wang, J. Zhang, L. Liu, X. Fan, C. Liu, M. Liu, L. Guo, P.Y. Lee, I. Aksentijevich, and Q. Zhou. 2023. A gain-of-function variation in PLCG1 causes a new immune dysregulation disease. J. Allergy Clin. Immunol. 152:1292–1302. doi:10.1016/j.jaci.2023.06.020.

Town, J.P., and O.D. Weiner. 2023. Local negative feedback of Rac activity at the leading edge underlies a pilot pseudopod-like program for amoeboid cell guidance. PLOS Biol. 21:e3002307. doi:10.1371/journal.pbio.3002307.

Tsai, F.-C., A. Seki, H.W. Yang, A. Hayer, S. Carrasco, S. Malmersjö, and T. Meyer. 2014. A polarized Ca2+, diacylglycerol and STIM1 signalling system regulates directed cell migration. Nat. Cell Biol. 16:133–144. doi:10.1038/ncb2906.

Tsien, R.Y. 1981. A non-disruptive technique for loading calcium buffers and indicators into cells. Nature. 290:527–528. doi:10.1038/290527a0.

Tvorogov, D., X.-J. Wang, R. Zent, and G. Carpenter. 2005. Integrin-dependent PLC-gamma1 phosphorylation mediates fibronectin-dependent adhesion. J. Cell Sci. 118:601–610. doi:10.1242/jcs.01643.

Waldo, G.L., T.K. Ricks, S.N. Hicks, M.L. Cheever, T. Kawano, K. Tsuboi, X. Wang, C. Montell, T. Kozasa, J. Sondek, and T.K. Harden. 2010. Kinetic scaffolding mediated by a phospholipase C–β and g_q_ signaling complex. Science. 330:974–980. doi:10.1126/science.1193438.

Waldron, R.T., and E. Rozengurt. 2003. Protein kinase C phosphorylates protein kinase D activation loop Ser744 and Ser748 and releases autoinhibition by the pleckstrin homology domain. J. Biol. Chem. 278:154–163. doi:10.1074/jbc.M208075200.

Wei, C., X. Wang, M. Chen, K. Ouyang, L.-S. Song, and H. Cheng. 2009. Calcium flickers steer cell migration. Nature. 457:901–905. doi:10.1038/nature07577.

Weiner, O.D., P.O. Neilsen, G.D. Prestwich, M.W. Kirschner, L.C. Cantley, and H.R. Bourne. 2002. A PtdInsP3- and Rho GTPase-mediated positive feedback loop regulates neutrophil polarity. Nat. Cell Biol. 4:509–513. doi:10.1038/ncb811.

Weiner, O.D., G. Servant, M.D. Welch, T.J. Mitchison, J.W. Sedat, and H.R. Bourne. 1999. Spatial control of actin polymerization during neutrophil chemotaxis. Nat. Cell Biol. 1:75–81. doi:10.1038/10042.

Welf, E.S., S. Ahmed, H.E. Johnson, A.T. Melvin, and J.M. Haugh. 2012. Migrating fibroblasts reorient directionality by a metastable, PI3K-dependent mechanism. J. Cell Biol. 197:105–114. doi:10.1083/jcb.201108152.

Welf, E.S., C.E. Miles, J. Huh, E. Sapoznik, J. Chi, M.K. Driscoll, T. Isogai, J. Noh, A.D. Weems, T. Pohlkamp, K. Dean, R. Fiolka, A. Mogilner, and G. Danuser. 2020. Actin-membrane release initiates cell protrusions. Dev. Cell. 55:723–736.e8. doi:10.1016/j.devcel.2020.11.024.

Wilkinson, H.N., and M.J. Hardman. 2020. Wound healing: cellular mechanisms and pathological outcomes. Open Biol. 10:200223. doi:10.1098/rsob.200223.

Wu, C., S.B. Asokan, M.E. Berginski, E.M. Haynes, N.E. Sharpless, J.D. Griffith, S.M. Gomez, and J.E. Bear. 2012. Arp2/3 is critical for lamellipodia and response to extracellular matrix cues but is dispensable for chemotaxis. Cell. 148:973–987. doi:10.1016/j.cell.2011.12.034.

Wu, Y.I., D. Frey, O.I. Lungu, A. Jaehrig, I. Schlichting, B. Kuhlman, and K.M. Hahn. 2009. A genetically encoded photoactivatable Rac controls the motility of living cells. Nature. 461:104–108. doi:10.1038/nature08241.

Yang, Y.R., J.H. Choi, J.-S. Chang, H.M. Kwon, H.-J. Jang, S.H. Ryu, and P.-G. Suh. 2012. Diverse cellular and physiological roles of phospholipase C-γ1. Adv. Biol. Regul. 52:138–151. doi:10.1016/j.advenzreg.2011.09.017.

Zeng, L., X. Zhang, Y. Xiong, K. Sato, N. Hajicek, J. Sondek, and X. Su. 2024. Hyperactive PLCG1 drives non-canonical signaling to promote cell survival. BioRxiv Prepr. Serv. Biol. 2024.12.17.628879. doi:10.1101/2024.12.17.628879.

Zhao, H., M. Hakala, and P. Lappalainen. 2010. ADF/cofilin binds phosphoinositides in a multivalent manner to act as a PIP(2)-density sensor. Biophys. J. 98:2327–2336. doi:10.1016/j.bpj.2010.01.046.

Zimmerman, S.P., R.A. Hallett, A.M. Bourke, J.E. Bear, M.J. Kennedy, and B. Kuhlman. 2016. Tuning the binding affinities and reversion kinetics of a light inducible dimer allows control of transmembrane protein localization. Biochemistry. 55:5264–5271. doi:10.1021/acs.biochem.6b00529.

